# Identifying Lethal Dependencies with HUGE Predictive Power from Large-Scale Functional Genomic Screens

**DOI:** 10.1101/2021.10.29.466419

**Authors:** Fernando Carazo, Edurne San José-Enériz, Marian Gimeno, Leire Garate, Estíbaliz Miranda, Carlos Castilla, Xabier Agirre, Ángel Rubio, Felipe Prósper

**Affiliations:** Departamento de Ingeniería Biomédica y Ciencias, TECNUN, Universidad de Navarra, San Sebastián, Spain; Programa Hemato-Oncología, Centro de Investigación Médica Aplicada, IDISNA, Universidad de Navarra, Pamplona, Spain; Centro de Investigación Biomédica en Red de Cáncer (CIBERONC), Madrid, Spain; Departamento de Hematología, Clínica Universidad de Navarra, Universidad de Navarra, Pamplona

## Abstract

Recent functional genomic screens -such as *CRISPR-Cas9* or RNAi screening-have fostered a new wave of targeted treatments based on the concept of synthetic lethality. These approaches identified LEthal Dependencies (LEDs) by estimating the effect of genetic events on cell viability. The multiple-hypothesis problem related to the large number of gene knockouts limits the statistical power of these studies. Here, we show that predictions of LEDs from functional screens can be dramatically improved by incorporating the “HUb effect in Genetic Essentiality” (HUGE) of gene alterations. We analyze three recent genome-wide loss-of-function screens - Project Score, CERES score and DEMETER score-identifying LEDs with 75 times larger statistical power than using state-of-the-art methods. HUGE shows an increased enrichment in a recent harmonized knowledgebase of clinical interpretations of somatic genomic variants in cancer (with an AUROC up to 0.87). Our approach is effective even in tumors with large genetic heterogeneity such as acute myeloid leukemia, where we identified LEDs not recalled by previous pipelines, including *FLT3-*mutant genotypes sensitive to *FLT3* inhibitors. Interestingly, *in-vitro* validations confirm lethal dependencies of either *NRAS* or *PTPN11* depending on the *NRAS* mutational status. HUGE will hopefully help discover novel genetic dependencies amenable for precision-targeted therapies in cancer.

## INTRODUCTION

The traditional concept of synthetic lethality consists of the concurrent loss of functionality of two genes resulting in cellular death. A relevant example is the effectiveness of PARP inhibitors in tumors with inactivated BRCA1 and BRCA2 [1]. In recent years, the advances in functional genomics triggered by large-scale loss-of-function screening -such as *CRISPR-Cas9* or RNA interference (RNAi) screens-have boosted the discovery of hundreds of novel targets and context-specific lethal dependencies(LEDs) [2–7], defined as any association between two genes that results in differential viability depending on their genetic context (**Supplementary Figure 1**).

Several studies have carried out large-scale functional genomic screens to identify genome-wide targets and LEDs[2–5]. The Project Score [4], the Achilles Project [5,6] and the Project DRIVE [7] are three studies that performed genome-wide gene-knockouts in cancer cells aiming at establishing novel targets and LEDs. The refinement of computational and technical tools have improved the potential of loss-of-function screening to identify cancer vulnerabilities [3,8,9]. However, the multiple testing problem, related to the large number of gene knockouts, limits the statistical power of these studies and, therefore, their potential to find new targets.

Here we show that previous efforts to predict LEDs from functional screening can be significantly improved by taking into account the “HUb effect” in Genetic Essentiality (HUGE) of some gene alterations: a few specific sets of gene alterations are statistically associated with large changes in the essentiality of multiple genes. These “hub” aberrations lead to more statistically reliable LEDs than other alterations that do not participate in such hubs. We incorporate the HUGE effect in the statistical analysis of three recent loss-of-function experiments of both The Project Score and The Achilles Project (two datasets) showing that the number of LEDs discovered for a given FDR considerably improves for both *CRISPR-Cas9* and RNAi screens.

Using acute myeloid leukemia (AML), breast cancer (BRCA), lung adenocarcinoma (LUAD) and colon adenocarcinoma (COAD) as disease models, we validate that the predictions are enriched in associations used in the clinic. Finally, we validated *in-vitro* an example of a therapy guideline based in LED selection in AML. The HUGE analysis will help discover novel tumor vulnerabilities in specific genetic contexts, providing valuable candidates -targets and genetic variants as biomarkers- for further personalized treatments in hematological diseases or other cancer disorders.

## RESULTS

### Lethal Dependency analysis reveals a hub effect in genetic essentiality as a key for increasing the predictive power in large-scale analyses of loss-of-function screens

One of the main statistical challenges to find LEDs by integrating genome-wide functional screens with -omics datasets is the multiple hypothesis testing problem. Correction for multiple hypothesis reduces the statistical significance of results (meaning a decreased detection rate and an increased false-positive rate). The Project Score presented a large-scale genome-wide *CRISPR-Cas9* screening analysis targeting 18,009 genes in 30 different cancer types, across 14 different tissues [4,10]. They presented a methodology to detect LEDs based on finding differences in genetic essentiality in cell lines associated with the presence of specific gene variants (ANOVA test [11] with the Storey-Tibshirani p-value correction). Following this procedure, the Project Score was able to identify genetic LEDs in 7 out of 14 individual tissues analyzed [4,10].

Analyzing Project Score’s data, we noticed that for each tumor type a few specific genetic alterations were significantly associated with the genetic essentiality of a large set of genes. This handful of genetic aberrations shows a hub effect, in which a gene variant is associated with large changes in the essentiality of multiple genes. We termed this behavior as “HUb effect in Genetic Essentiality” (HUGE) (**Figure 1a**; other tumor types can be visualized in https://fcarazo.shinyapps.io/visnetShiny/). From the point of view of statistics, the HUGE effect is defined as improvement of the the statistical power by using gene variants as co-variates in a multiple hypothesis problem. Other biological covariates such as gene expression or copy number alterations has also shown to be covariates that increase the statistical power [12]. Using gene variants as statistical covariates provides a larger number of positives for a given FDR, which consequently means an increased specificity and sensitivity, or type I and type II errors, as demonstrated in **Supplementary methods, Section 6**. Interestingly, the analysis shows that HUGE effect is present in all tumors analyzed, significantly improving the predictive power of LEDs.

**Figure 1.**
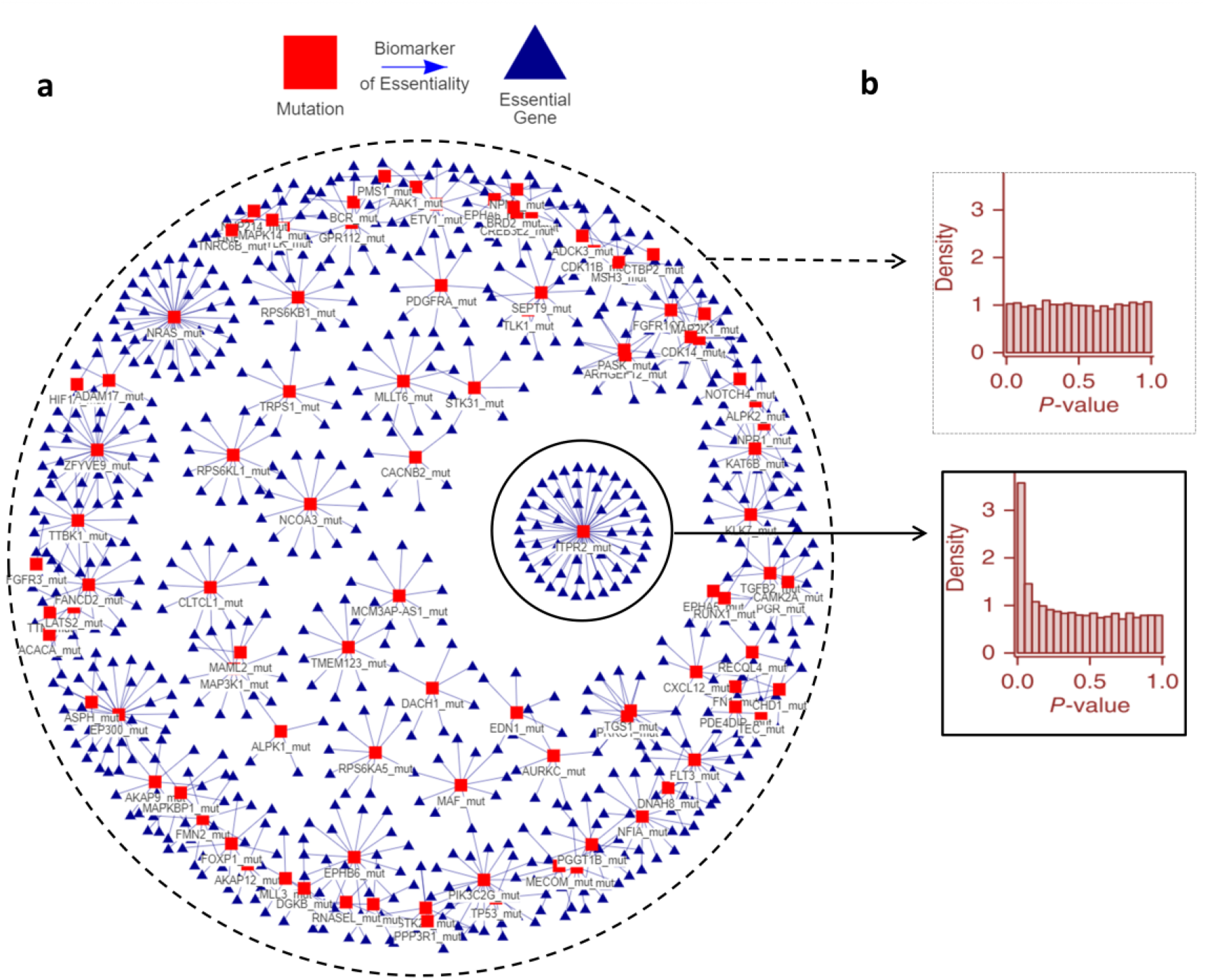
The hub effect in genetic essentiality (HUGE) in Acute Myeloid Leukemia: in a given cell, a small set of gene aberrations is associated with large changes in genetic essentiality. (a) A bipartite graph in which red squares represent gene variants (e.g., mutations), blue triangles represent significant changes in cell viability related to knocked-down genes. Both nodes are linked by a line if the variations in the essentiality have a statistically significant association with the presence of the gene variant. (b) Implications in p-value histograms of the HUGE effect. Hub associations show a high peak close to zero p-values indicating that the null hypothesis is rejected in more cases and that these genetic variants are associated to a higher response to the inhibition of more gene products. Segregating the statistical analysis according to the alteration provides more statistical power. Essential genes and other tumor types can be visualized in https://fcarazo.shinyapps.io/visnetShiny/.

The presence of the HUGE effect in a cancer type can be also understood as a predictive model in which each mutation has a different capability to define the genetic essentiality of multiple genes. To show it visually, the histogram of p-values of a gene alteration represents how gene alterations are associated with the genetic essentiality of multiple genes. Histograms of the p.values for alterations that conform to a “hub” show a peak near the origin, which means that cells with these alterations are sensitive to the depletion of a large number of genes (**Figure 1b**).

Conversely, if the hubs of alterations are not taken into account, the relationships of mutations and viability show a flat histogram of p-values. This does not necessarily mean that such relationships are not biologically relevant, but that is difficult to distinguish them from random associations, and will be considered as artifacts after multiple testing correction.

The HUGE effect helps palliate the multiple hypothesis correction problem. Using the mutation under study as a covariate, multiple hypotheses can be differently treated taking into account the overall association of gene alteration in the complete set of essential genes (**Supplementary Figure 2 and 3**). Using this concept, we developed a statistical model that integrates HUGE information to find LEDs (**Supplementary Figure 2**).

Previous efforts to correct multiple testing in this problem consider a single set of tests (all gene aberrations and *CRISPR-Cas9* knockouts) and apply a correction that controls the FDR, such as Storey-Tibshirani (ST), as done in the Project Score. Interestingly, in all tumors our approach increases the statistical power of the analysis. From a statistical point view, a flat histogram is compatible with the null hypothesis for all the tests and, therefore, multiple hypothesis correction drives to none or few discoveries (**Supplementary Figure 4**). Every single tumor show p-value histograms related to specific gene variants that have a higher zero-peak than the histogram associated to all tests in such tumor (**Supplementary Figures 5-23**).

To test this approach, we compare the results using HUGE with previous LED identification strategies in three genome-wide functional genomic projects: Project score, the DEMETER score and the CERES score (DEMETER and CERES are included in the Achilles Project). First, to test the potential of HUGE to predict LEDs with *CRISPR-Cas9* screens, we analyze the Project Score dataset [4]. Project Score integrates 215 different genetic events across 14 tumor types, including SNVs and CNVs. In the same reference, the authors found at least one LED in 7 out of the 14 tumor types analyzed. 40 out of 215 events were detected to be significant biomarkers of essentiality (FDR ≤ 20%), which correspond to 77 unique LEDs (a single genetic event can be associated with several essential genes). Analyzing Project Score’s data using the HUGE-based methodology, we identify 1,438 unique associations with the same FDR (18 times larger than Project Score, **Figure 2a**), corresponding to 80 single genetic events. Besides, using HUGE we detect at least one LED in all the 14 tumors analyzed, finding LEDs in 10 tumors that would have been missed using the original pipeline, affecting around 10-20 genes for each disease type.

**Figure 2.**
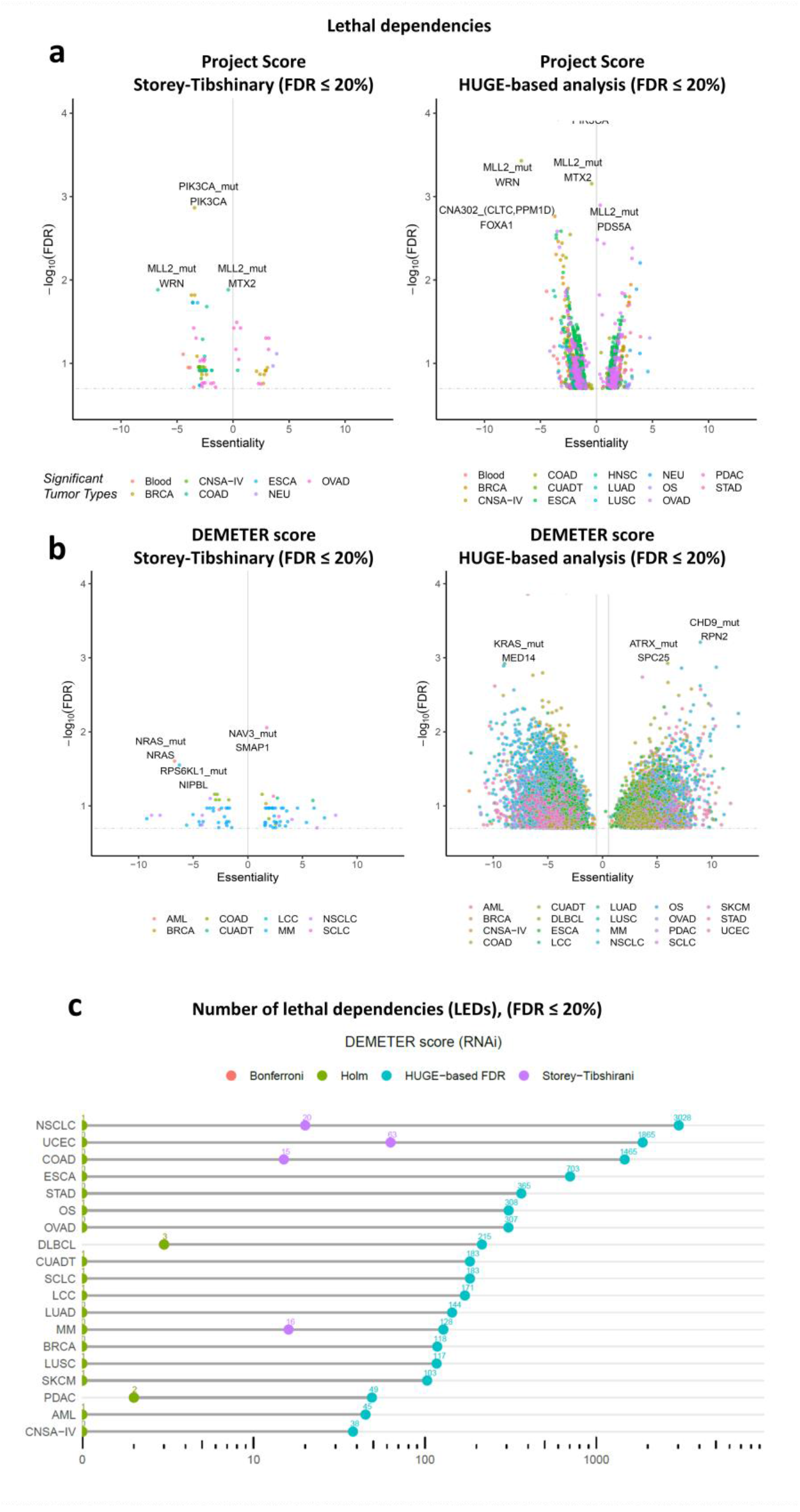
HUGE-based analysis with Project Score and Achilles Project datasets. (a) Volcano plots of lethal dependencies (LEDs) identified in the Project Score dataset. From left to right: i) result of Project Score, ii) results of analyzing Project Score dataset with HUGE-based methodology. Each dot represents a significant LED (FDR<20%). The X-axis represents the difference in gene essentiality when the event (gene variants) is present. The Y-axis represents the FDR values (−log10) for that change. (b) Equivalent volcano-plots using Achilles Project. From left to right: i) results of Achilles Project analyzed with the standard procedure, ii) results of analyzing Achilles Project dataset with HUGE-based methodology. (c) The number of LEDs found (FDR ≤ 20%) in 19 tumors of the DEMETER score (RNAi) and 22 tumors of the CERES score (CRISPR-Cas9) using standard statistical pipelines (Storey-Tibshinary, Bonferroni, and Holm) and the HUGE-based algorithm. Bonferroni and Holm return the same number of hypotheses in all cases. LEGEND: ALL: acute lymphoblastic leukemia; AML: acute myeloid leukemia; BRCA: breast ductal carcinoma; CNSA-IV: central nervous system astrocytoma grade IV; COAD: colon adenocarcinoma; CUADT: upper aero-digestive tract squamous cell carcinoma; DLBCL: diffuse large B-cell lymphoma; ESCA: esophagus squamous cell carcinoma; KIRC: kidney renal clear cell carcinoma; LCC: lung large cell carcinoma; LUAD: lung adenocarcinoma; LUSC: lung squamous cell carcinoma; MM: multiple myeloma; NSCLC: non–small cell lung carcinoma; OS: osteosarcoma; OVAD: ovary adenocarcinoma; PDAC: pancreas ductal carcinoma; SCLC: small cell lung carcinoma; SKCM: skin carcinoma; UCEC: endometrium adenocarcinoma.

We also tested HUGE in the DEMETER score of the Achilles Project to predict LEDs, in this case using RNAi screening. The DEMETER dataset [5,13] is a large-scale genome-wide experiment of RNA interference libraries (17,085 knockdown genes) in 19 tumor types (**Supplementary Table 5**). We integrate the DEMETER data with the corresponding cell line gene alteration profiles (genetic variants in ^~^1,600 genes) obtained from the Cancer Cell Line Encyclopedia (CCLE) [14] and Shao et al. [6]. This integration turns out to have 27 Million hypotheses, which will hardly impair p-values after multiple hypothesis correction (**Supplementary Figure 2**). Then, we replicate the Project Score’s pipeline with the DEMETER dataset and compare it with the HUGE-based approach to find LEDs, also including in the comparison other two standard p-value corrections used to control the FDR, namely Holm and Bonferroni. Using the standard ST procedure, we find 126 LEDs (FDR ≤ 20%). There are LEDs for 7 out of 19 tumors. The same dataset and FDR threshold using the HUGE-based approach provides 9,535 LEDs (75.7 times larger than using ST). All cancer types (19 out of 19) showed significant LEDs in the HUGE-based analysis (**Figure 2b**). HUGE identifies 1,675 LEDs in 6 tumor types in which other methods recall no LEDs (FDR ≤ 20%); and 9,409 LEDs in 19 tumor types that would have been missed using previous procedures (FDR ≤ 20%; **Figure 2c**). These results show that the HUGE effect is present with different intensities in all tumor types analyzed (**Supplementary Figures 5-23**).

As a further test of the increase predictive power of HUGE we carry out a similar analysis using the CERES score, a *CRISPR-Cas9* experiment of 22 tumors also included in the Achilles Project. In this case, the number of significant pairs is enriched 14 times over the standard approaches (FDR ≤ 20%; **Supplementary Figure 24**).

### LEDs predicted by HUGE have better validation rates than standard approaches

Validating a ranking of LEDs is not a simple task: it is desirable to have a gold standard of disease-specific list of validated target-biomarker associations. We select as our gold standard The Variant Interpretation for Cancer Consortium (VICC) Meta-Knowledgebase [15,16]. This database integrates different datasets of clinical associations and includes the level of evidence for each entry: spanning from professional FDA guidelines to preclinical findings.

We test the enrichment in associations included in VICC in four tumor types, namely acute myeloid leukemia (AML), breast cancer (BRCA), lung adenocarcinoma (LUAD) and colon adenocarcinoma (COAD) for both HUGE and standard statistical methods. The VICC knowledgebase integrates (in September 2021) 19,551 clinical interpretations of somatic genomic variants in cancer of both resistant and sensitive biomarkers. We delete duplicated and incomplete associations, focused on those related to confirmed mutations and manually selected associations that match each tumor type (including synonyms).

We first run the two procedures (HUGE and Storey-Tibshirani; ST) with AML cell lines (**Supplementary Table 6**) to find LEDs and compare how many LEDs predicted by HUGE and by ST are included in the VICC knowledgebase. For instance, if HUGE or the ST procedure predicts *FLT3* mutant AML genotypes to be sensitive to *FLT3* inhibition, it will be considered a true positive LED, as FLT3 is a well-known target of AML and mutations in FLT3 are known to be sensitive biomarkers of the effectiveness of most FLT3-inhibitors [17,18].

In total, 216 out of 19,551 associations matched these filters. Getting the top 500 LEDs according to the ranking using the HUGE algorithm with AML, we find 17 LEDs that match the VICC knowledgebase of known clinic relationships (**Supplementary Table 1**; Fisher p-value < 1e-51). An equivalent analysis using the standard pipeline (ANOVA test [11] with the Storey-Tibshirani p-value correction) shows that out of the top 500 LEDs, only 1 is included in the VICC knowledgebase (**Supplementary Table 2;** Fisher p-value = 6.551e-3). This means that HUGE analysis identifies 16 true positive dependencies not recovered by ST (Fisher p-value = 6.41e-5). The global value of AUROC (0.53) is not too far from the baseline of 0.5 (**Figure 3a**), perhaps because of the scarcity of true positives in our gold standard. We perform the same analysis with LUAD, BRCA and COAD getting AUCROC values of 0.62 (vs 0.5), 0.87 (vs 0.64) and 0.72 (vs 0.54) for HUGE and ST respectively. All cases show better values for HUGE than for ST (**Figure 3b-d, Supplementary Figure 25**).

**Figure 3.**
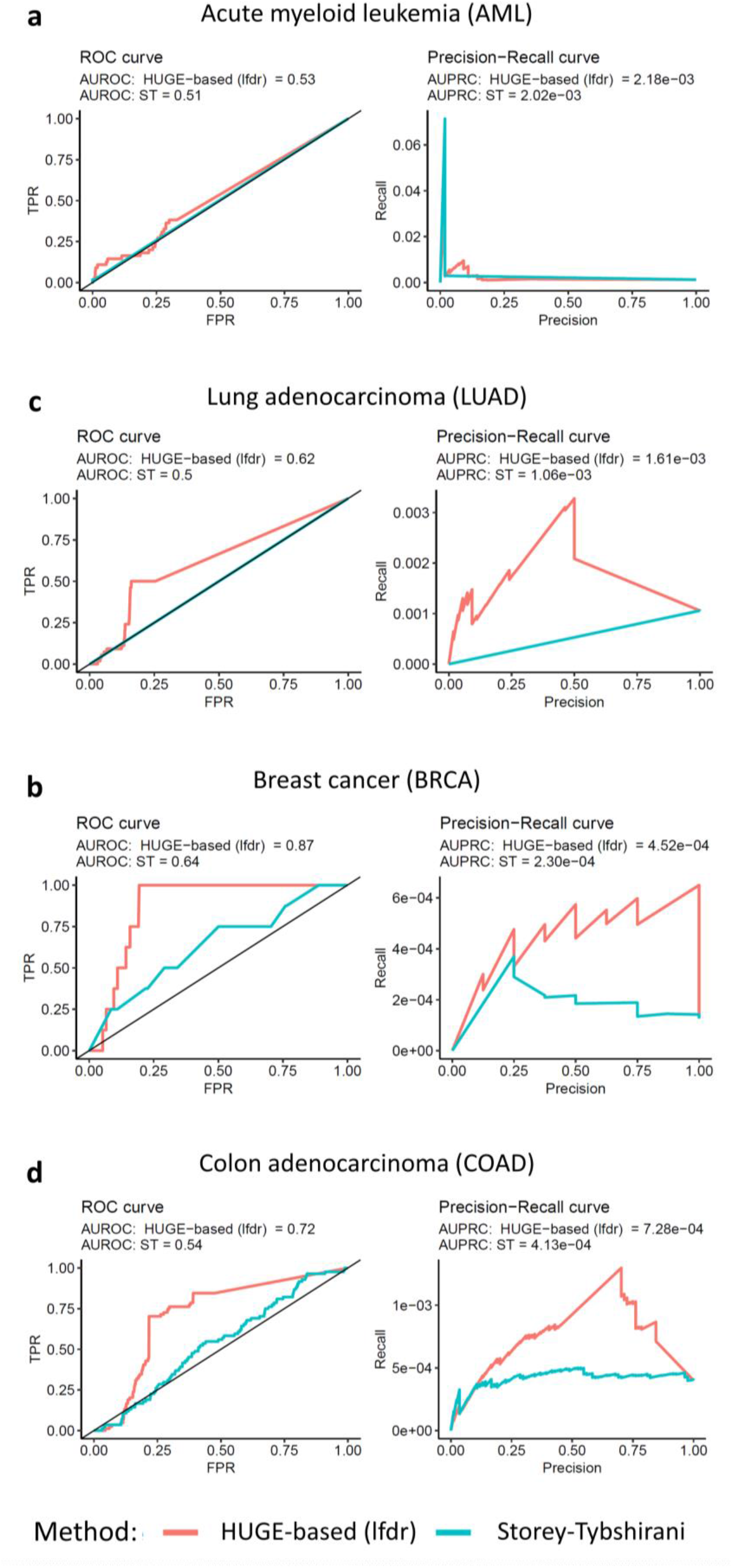
ROC and precision-recall curves of four tumor types. True positives were found. somatic genomic variants in the knowledgebase of the Variant Interpretation for Cancer Consortium (VICC). a) AML, b) BRCA, c) LUAD and d) COAD. We selected associations indicated for each tumor type that are within the three highest levels of confidence (Level A: Evidence from professional guidelines or FDA-approved therapies relating to a biomarker and disease; Level B: Evidence from clinical trials or other well-powered studies in clinical populations, with expert consensus; and Level C: Evidence for therapeutic predictive markers from case studies, or other biomarkers from several small studies, or evidence for biomarker therapeutic predictions for established drugs for different indications).

### Applying HUGE methodology to acute myeloid leukemia cell-lines discovers potential therapy biomarkers

AML is a hematologic neoplasm characterized by a remarkable phenotypic and genomic heterogeneity [19], a challenging disease model to test the applicability and impact of HUGE. We run the complete HUGE pipeline with AML and validate *in vitro* two of the predicted LEDs.

As a preliminary step, we identify the potential genes that are essential for AML cell survival. The Achilles Project yield 443 essential genes that are essential and specific for AML cells compared to other tumors (**Supplementary Table 3**). Some of these genes belong to pathways known to be deregulated in AML (e.g., MYB [20] or CEBPA [21]). Interestingly, 160 of these 443 genes have previously been identified as potential cancer drivers in hematological malignancies according to the Candidate Cancer Gene Database (p-value = 7.76e-05, Fisher exact test) [22].

We then run the HUGE algorithm to identify genomic alterations that could be defined as LED partners of those 443 essential genes. In this pipeline, we require predicted pairs to be biologically related to each other in the STRING database (see **Online Methods**). LED associations can be broken down into three groups regarding their dependency type: *positive lethal dependency (pLED)*, when a gene variant marks sensitivity to the inhibition of another gene; *negative lethal dependency (nLED)l*, when a gene variant marks resistance to the inhibition of another gene; or *dual lethal dependency (dLED)*, when the same gene variant confers, concurrently, sensitivity to the inhibition of one gene and resistance to the inhibition of another gene (**Supplementary Figure 1**). In total, we predict 24 LEDs, (12 *pLEDs* and 12 *nLEDs*, including 2 *dLEDs;* p-value < 0.05, local FDR ≤ 0.6 and |ΔEssentiality| > 2; **Figure 4a**, **Table 1, Supplementary Figure 26**, and **Supplementary Table 4**). Using the standard multiple hypotheses correction only 1 dependency turns out to be statistically significant. We provide the identified LEDs for the 19 tumors included in the Achilles Project following a similar pipeline (**Supplementary Tables 8 to 26**).

**Figure 4.**
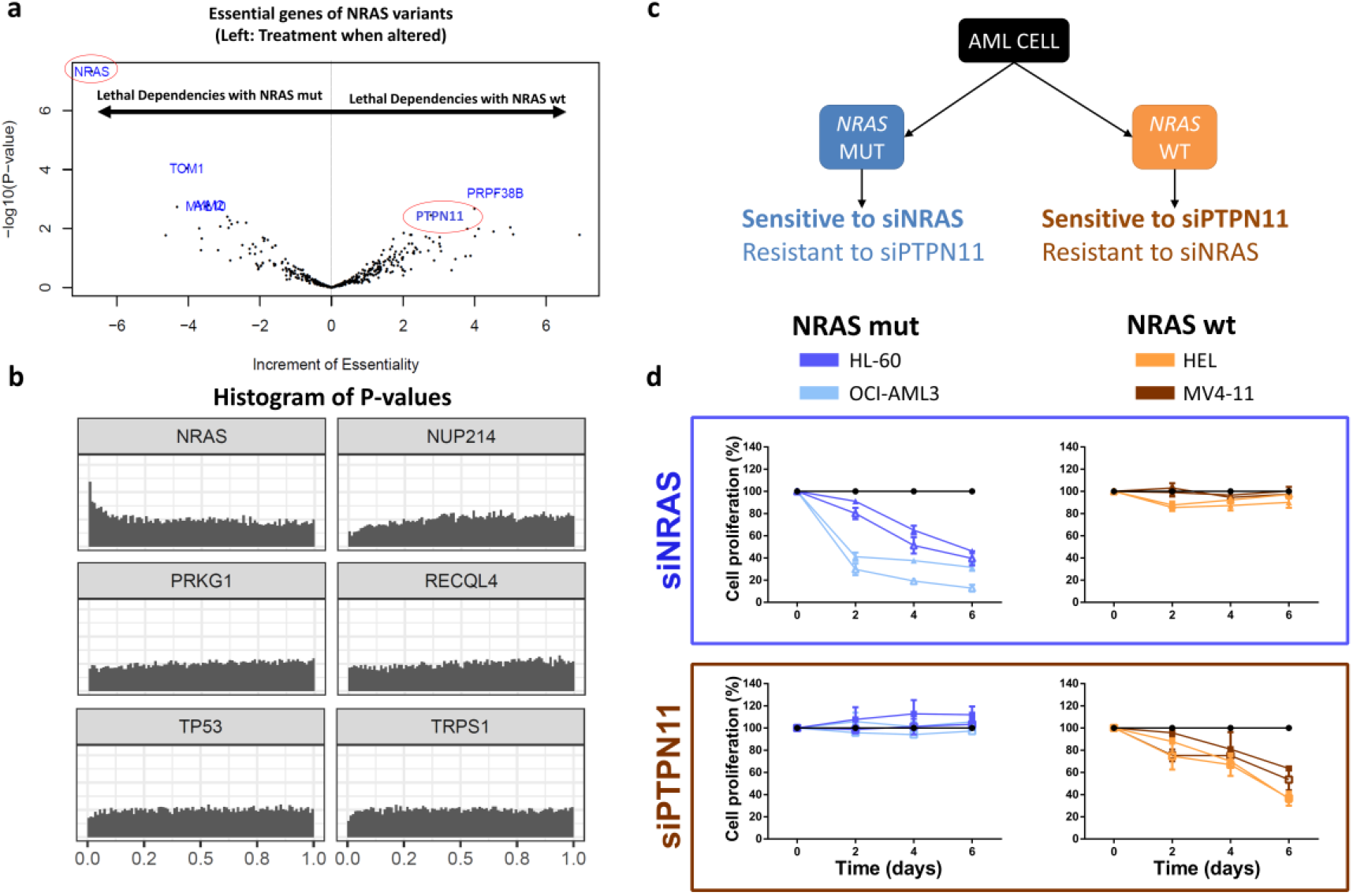
Gene variants-based treatment guidelines in acute myeloid leukemia (AML). (a) Volcano-plot of LEDs related to NRAS genetic mutations (left; MUT) and -wildtype (right; WT) phenotypes. Increment of Essentiality and −log10 (p-value) are shown in X-axis and Y-axis, respectively. (b) Histogram of p-values for 6 genetic sequence variants in AML. NRAS-alteration is enriched in close to zero p-values, which is the basic concept of HUGE-based statistical approach. All genetic variants histograms of p-values can be found in the Supplementary Material. (c) Summary of the computational predictions validated: NRAS-altered cells were predicted to be sensitive to siNRAS and resistant to siPTPN11. Conversely, NRAS-wt cells were predicted to be sensitive to siPTPN11 and resistant to siNRAS. (D) Tumor proliferation of the four AML cell lines after inhibiting NRAS (siNRAS) and PTPN11 (siPTPN11) with specific siRNAs. Blue: NRAS-altered AML cell lines (HL-60 and OCI-AML3); Orange: NRAS-wild-type AML cell lines (MV4-11 and HEL).

**Table 1.** Ranking of generalized synthetic lethality in AML using the HUGE-based statistical approach. The ranking is divided into three groups regarding the typology of the lethal dependency relationship: Positive Synthetic Lethality, Negative Synthetic Lethality or Dual Synthetic Lethality (**Figure 1**). The Increment of Essentiality column represents the average variation in the DEMETER score between altered and wild-type cells, and its sign is related to the lethal depency relationship. Lethal dependencies that share the same essential gene and the same Increment of Essentiality sign have been omitted in this table (see complete data in **Supplementary Table 2**).

| Gene variant<br>Biomarker | Essential<br>Gene | Increment of<br>Essentiality | t-score | p value | Local FDR |
| --- | --- | --- | --- | --- | --- |
| **Positive Lethal Dependencies** |  |  |  |  |  |
| TGS1 | SNRPF | -7,87 | -4,05 | 6,69E-04 | 3,36E-01 |
| CLTCL1 | UBR5 | -6,66 | -3,59 | 1,99E-03 | 2,20E-01 |
| FLT3 | FLT3 | -6,36 | -4,53 | 2,28E-04 | 2,00E-01 |
| CDK14 | CDK2 | -3,95 | -2,75 | 1,28E-02 | 4,30E-01 |
| AURKC | ACTL6A | -3,26 | -3,89 | 9,55E-04 | 4,99E-01 |
| **Negative Lethal Dependencies** |  |  |  |  |  |
| NPM1 | EEF2 | 3,81 | 3,34 | 3,39E-03 | 5,96E-01 |
| PIK3C2G | CDK6 | 3,35 | 2,95 | 8,20E-03 | 3,51E-01 |
| NCOA3 | EP300 | 3,04 | 2,75 | 1,25E-02 | 4,94E-01 |
| CDK14 | CCND2 | 2,97 | 2,22 | 3,88E-02 | 4,99E-01 |
| EPHB6 | ZNF266 | 2,53 | 2,77 | 1,22E-02 | 3,42E-01 |
| ZFYVE9 | TOM1L2 | 2,14 | 2,35 | 2,96E-02 | 5,12E-01 |
| **Dual Lethal Dependencies** |  |  |  |  |  |
| <u>NRAS</u> | <u>NRAS</u> | <u>-6,83</u> | -8,71 | 4,67E-08 | 1,38E-04 |
| <u>NRAS</u> | <u>PTPN11</u> | <u>4,17</u> | 2,2 | 4,05E-02 | 5,89E-01 |
| EP300 | PLK1 | -8,11 | -4,04 | 7,01E-04 | 2,17E-01 |
| EP300 | KLF2 | 3,69 | 4,08 | 6,38E-04 | 2,12E-01 |

*NRAS* mutation ranks first in the analysis. Lethally dependent partners associated with *NRAS* genetic sequence variants show a p-value histogram that peaks at the origin (**Figure 4a and 4b**), meaning that *NRAS* mutations are associated with more tumor vulnerabilities than other alterations. Interestingly, *NRAS* alteration forms a *Dual Lethal Dependency* with *PTPN11* (**Table 1, Figure 4c**): it confers tumor sensitivity to *NRAS* inhibition and resistance to *PTPN11* inhibition.

To validate our prediction, we first check that both *NRAS* and *PTPN11* siRNAs efficiently decreased the *NRAS* and *PTPN11* expression, respectively, in four AML cell lines (**Supplementary Figure 27**). Then, we confirm the computational hypothesis: the downregulation of *NRAS* significantly decreases cell proliferation only in the *NRAS*-altered AML cell lines, and the inhibition of *PTPN11* expression produces an equivalent effect, but specifically in the *NRAS*-wt AML cell lines (**Figure 4d**), validating the predicted dLED. Remarkably, the validated *PTPN11-NRAS*-wt pair was not detected using standard methodologies.

## DISCUSSION

The advent of large-scale functional genomic screens has allowed the identification of hundreds of novel gene targets and the prediction of genome-wide LEDs [4,23]. This strategy has multiplied treatment strategies, as using LEDs, the drug targets can be decoupled from their corresponding predictive biomarkers. The main statistical limit to find LEDs is the large number of hypotheses that result from integrating gene essentiality and genetic functional events. In this work, we present HUGE, a novel analysis of *CRISPR-Cas9* and RNAi large-scale screens that significantly improves the predictive power to find LEDs from loss-of-function screens in human tumors. It relies on the fact that some gene alterations are statistically related to the essentiality of large sets of genes. Using this characteristic as a prior covariate we significantly improve the predictive power of LEDs.

Notably, the presence of the HUGE effect does not necessarily mean biological causality. HUGE dependencies are more statistically reliable than others, but this does not imply that predicted alterations are the major players on tumor development; i.e., they can be just genetic biomarkers of gene essentiality. However, the fact that gene alterations co-occur with multiple LEDs in genetic hubs can be exploited to improve the statistical power.

To measure the increased predictive power of HUGE, we carry out three different comparisons within three functional genomic datasets: the Project Score, the DEMETER score and the CERES score. HUGE identifies LEDs with 14 and 75 times larger statistical power than using state-of-the-art methods in *CRISPR-Cas9* and RNAi, respectively. However, it could be argued that this result could be an artifact of the statistical technique and that lowering the threshold for standard procedures would provide LEDs with similar reliability. This is not the case. As shown in the results, using the same number of predictions, HUGE’s results are more enriched in clinically used biomarkers than ST’s results. Remarkably, one of the 16 LEDs only identified by HUGE is the known interaction of *FLT3-*mutant genotypes sensitive to *FLT3* inhibitors, such as Midostaurin. This fact is only an example of the key importance of considering the HUGE effect when analyzing LEDs with large-scale functional screens.

A p-value histogram can be modeled as the superposition of two distributions, a uniform distribution (which corresponds to the null hypothesis) and another distribution with a larger proportion of low p-values. A good covariate splits the overall p-value histogram into histograms with different enrichments in small p-values. If all the histograms related to a covariate have similar shapes, it means that the covariate is uninformative. Here, we show that stating which gene is mutated in each test is a good covariate for the LED prediction problem because there is a hub effect of gene aberrations in gene essentiality. The usage of covariates has successfully been incorporated before in other genomics applications (e.g., the abundance of a gene is known to be informative in differential expression analyses; or proximity of loci in the genome is known to play a role in genome-wide association studies), but it has not yet been exploited in large-scale functional genomic screens.

Predicting true LEDs is especially challenging for tumors with high genetic heterogeneity. In AML, for instance, state-of-the-art approaches only recover 2 LEDs. The HUGE-based approach captured 24 LEDs for the same False Discovery Rate (FDR). Interestingly, *NRASwt-PTPN11* LED, which was only identified by HUGE, has been validated *in vitro*. The validation in AML highlights the potential of the HUGE-based approach to discover and validate new LEDs of biomarkers and drug targets. We pinpoint the *duLED* characteristic of the *NRAS* gene, meaning that if a tumor has *NRAS* mutated a treatment that targets *NRAS* itself would be the best option to reduce their tumorigenicity, whereas if it is *NRAS* wild-type, a *PTPN11* inhibition would be a better recommendation. This methodology has potential applications both in basic and clinical research.

In conclusion, this work provides a computational approach to identify LEDs with increased predictive power. This analysis opens new possibilities to use genetic variants as predictive events for precision oncology, by analyzing both previous and future functional genomic screens. Besides, this analysis enhances current applications in translational oncology, such as drug development or drug repositioning projects.

## ONLINE METHODS

### Data integration

Data of loss-of-function screens libraries (17,980 knockout genes in 412 cancer cell lines) of the project Achilles [13] were integrated with gene expression and their corresponding gene alteration profiles (gene variants in ^~^1600 genes; **Figure 2a**) obtained from CCLE and Shao et al. [6]. We gathered gene expression of cells using RNA-seq data to confirm that the genes that were essential for a cohort of cells were expressed before the RNAi library experiment was performed [24]. Gene variant panels were filtered out using the parameters of CCLE’s authors to avoid common polymorphisms, low allelic fraction, putative neutral variants, and substitutions located outside of the coding sequence [25].

We used the DEMETER score [5,8] as a measure of gene essentiality of the RNAi libraries of the project Achilles [13]. DEMETER quantizes the competitive proliferation of the cell lines controlling the effect of off-target hybridizations of siRNAs by solving a complex optimization problem. The more negative the DEMETER score is, the more essential the gene is for a cell line. We imputed missing elements of DEMETER using the nearest neighbor averaging algorithm [26]. Besides, we collected gene expression patterns from RNA-seq data [24] to confirm that essential genes are expressed when they are essential. Based on DEMETER data, we first identified genes that were essential for a selected tumor subtype. Essential genes were required to meet several criteria: i) they must be essential for at least 20% samples of the selected cancer subtype, ii) they must be specific to the cancer type under study, i.e. they must be non-essential for other cancer types and iii) they must be expressed before RNAi experiment (>1TPM at least in 75% samples).

### Statistical model

We developed a statistical algorithm to identify genes whose essentiality is highly associated with the genetic alteration of other genes. Dealing with this statistical issue implies solving a large multiple hypotheses problem (more than one million hypotheses). In similar scenarios, traditional corrections -such as Benjamini-Hochberg (BH), Bonferroni, or Holm-showed very few or no gene-biomarker LEDs for a given FDR [12]. To overcome this problem, we developed a covariate-based statistical approach -similar to the Independent Hypothesis Weighting procedure [12] (**Supplementary Figure 2)**.

Let *e* denote the number of RNAi target genes and *n* denote the number of screened samples. Let **D** be an *e* × *n* matrix of essentiality whose entries d_ij_ represent the DEMETER score for the RNAi target *i* in sample *j*. Let **m** be a *m*×*n* dichotomized matrix whose entry *m_ij_* denotes whether sample *j* is mutant or not according to the previous criteria:

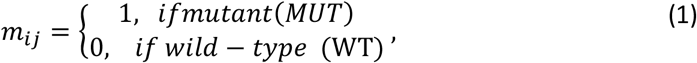

Let **s** be a subset of n’ cell lines that yields an essentiality vector**d**_s_ =(*d_e_s_1___*,…,*d_e_s_n___*) for the e^th^ RNAi target. Let ***m**_s_* = (*m_s_1__*…, *m_S_n__*) be the expression vector of a putative gene biomarker. The null hypotheses are defined as:

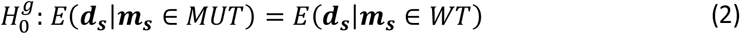

This null hypothesis is, therefore: “the expected essentiality of a gene knock-down is identical in mutant and wild-type cell lines”. To test this hypothesis, we used a moderated t-test implemented in *limma*[27]. We applied this test for each RNAi target and all the gene variants to get the corresponding p-values (**Supplementary Figure 2**). Dealing with these p-values implies correcting for multiple hypotheses.

In our case, we divided the p-values corresponding to all the tests into n groups, where n is the number of altered genes. For each of these groups, we computed the local false discovery rate (local FDR) [28]. The local FDR estimates, for each test, the probability of the null hypothesis to be true, conditioned on the observed p-values. The formula of the local FDR is the following:

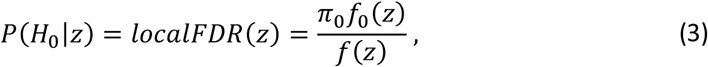

where *z* is the observed p-values, Π_0_ is the proportion of true null hypotheses –estimated from the data-, *f*_0_(*z*) the empirical null distribution –usually a uniform (0,1) distribution for well-designed tests- and *f*(*z*) the mixture of the densities of the null and alternative hypotheses, which is also estimated from the data.

As stated by B. Efron and R. Tibshirani [29], “the advantage of the local FDR is its specificity: it provides a measure of belief in gene i’s ‘significance’ that depends on its p-value, not on its inclusion in a larger set of possible values” as it occurs, for example with q-values or the standard FDR. The local FDR and Π_0_ were estimated using the Bioconductor’s R Package *qvalue* [30].

### Comparison with the Project Score

To compare our results with Project Score’s ones, we selected the same 12 primary cancer tissues shared in both datasets. The comparison followed two steps: 1) using CCLE and DEMETER scores with the Project Score’s algorithm, 2) running our approach adapted to Project Score conditions. In the first step, following the code published in their work, an ANOVA test was performed on each tissue to calculate all possible dependent partners. The Storey-Tibshirani correction was then used, using the criteria mentioned in Project Score methods [4]. This enabled us to correct the ANOVA *p*-values and get significant associations. Secondly, the comparison between both methodologies was only possible if the same adjusted p-value is calculated for both datasets. Therefore, we estimated the FDR with our data as it is the q-value selected by the Project Score. The FDR correction was obtained using the Bioconductor R package *IHW* [12], which enables the consideration of covariates-based multiple hypothesis correction, as well as estimating the FDR. Discoveries from both methodologies in DEMETER and CCLE datasets were plotted in different volcano plots, and the number of significant LEDs were counted (FDR<20%).

### Integration of the VICC knowledgebase of clinical interpretations of genomic variants

We downloaded 19,551 clinical interpretations of somatic genomic variants in cancer of the Variant Interpretation for Cancer Consortium (VICC) [15,16] (version December 2020). We filtered out incomplete (e.g., entrees without annotated drug or biomarker) and redundant associations. We then selected all associations that are annotated to acute myeloid leukemia (AML) and synonyms. From all drugs, we selected those that have an annotated protein target. To do so, we retrieved the data publicly available in the ChEMBL [31] and DrugBank [32] online repositories. In total, 216 out of 19,551 associations matched these criteria. We consider a true positive if either HUGE or ST identifies an LED whose mutation biomarker coincides with a VICC’s association and the protein target is included in the same association, or at least in a gene of the same pathway in the STRING database (v.11, STRING score threshold = 400; default value on STRING for “medium” confidence) [33].

We calculated ROC and PR curves considering the two top evidence levels included in VICC [15,16], namely, (i) evidence from professional guidelines or FDA-approved therapies; and (ii) evidence from clinical trials or other well-powered studies in clinical populations, with expert consensus.

### Application to acute myeloid leukemia as a disease model

We applied the pipeline to the AML cohort of cell-lines (n=15). In the first step, essential genes were required to be: (i) essential for at least 25% AML samples, (ii) specific for AML cells, and (iii) expressed before the RNAi experiment. The algorithm outputs a ranking of significant gene pairs (LEDs) that consist of a couple of genes in which the first one is essential depending on the genetic alteration of the other.

For the final ranking for AML, we selected those LEDs that showed a p-value < 0.05 and local FDR ≤ 0.6, |ΔDEMETER| > 2 (default value suggested by DEMETER’s authors). Additionally, we interrogated which of these LEDs had direct relationships (co-expressed, annotated in the same pathway database, or contained in a common experiment) in the STRING database [33] to ensure there is an established biological relationship between the essential gene and the subrogate biomarker. This biological double-check is not necessary and can be omitted when the researcher looks for novel relationships.

*In vitro* validation was performed using siRNAs against *NRAS* and *PTPN11* in four different AML cell lines, two with *NRAS*-genetic variants (HL-60 and OCI-AML3) and two *NRAS*-wt cell lines (MV4-11 and HEL). Finally, the model was compared with 3 standard statistical methods (namely Benjamini-Hochberg (BH), Bonferroni and Holm) known to have suboptimal sensitivity (recall of true positives) in specific scenarios in 19 additional tumor subtypes to define the potential for controlling the FDR. [12] See Supplemental Methods for more details.

## Acknowledgments

This research was funded by the Provincial Council of Gipuzkoa through the MINEDRUG project, by grants from Instituto de Salud Carlos III (ISCIII) PI14/01867, PI16/02024, PI17/00701, PI19/01352, PI20/01306 and TRANSCAN EPICA AC16/00041 (Co-financed with European Union FEDER funds), Fundació La Marató de TV3, Cancer Research UK [C355/A26819] and FC AECC and AIRC under the Accelerator Award Programme, CIBERONC CB16/12/00489 (Co-financed with European Union FEDER funds), Spanish Ministry of Economy, Industry and Competitivity (RTHALMY SAF2017-92632-EXP), Gobierno de Navarra, Departamento de Salud 40/2016 and Departamento de Industria (Proyecto Estrategico, Reto Genomica, DIANA). FC was partially supported by a Basque Government predoctoral Grant [PRE_2016_1_0194]. The authors would like to thank Francisco J. Planes, Luis V. Valcárcel, Xabier Cendoya and Lucía Campuzano for their fruitful comments on the development of the methodology.

## Authorship contribution

Conception and design: FC, ESJ-E, XA, AR and FP. Development of statistical model: FC, MG and AR. Acquisition of data (provided cell lines, provided siRNAs, etc.): ESJ-E, LG, EM, XA and FP. Carrying out experiments: ESJ-E, LG, EM and XA. Analysis and interpretation of data (e.g., statistical analysis, biostatistics, computational analysis): FC, ESJ-E, MG, CC, XA, AR and FP. Writing, review, and/or revision of the manuscript: FC, ESJ-E, MG, XA, AR and FP. Administrative, technical, or material support (i.e., reporting or organizing data, constructing vignettes): FC and AR. Study supervision XA, AR and FP. All authors read and approved the final manuscript.

## Ethical Approval

The AML cell lines used in this study were purchased from ATCC or DSMZ and were authenticated by performing a short tandem repeat allele profile.

