## Supplementary Material for "Identifying Lethal Dependencies with HUGE Predictive Power from Large-Scale Functional Genomic Screens"

### SUPPLEMENTARY FIGURES

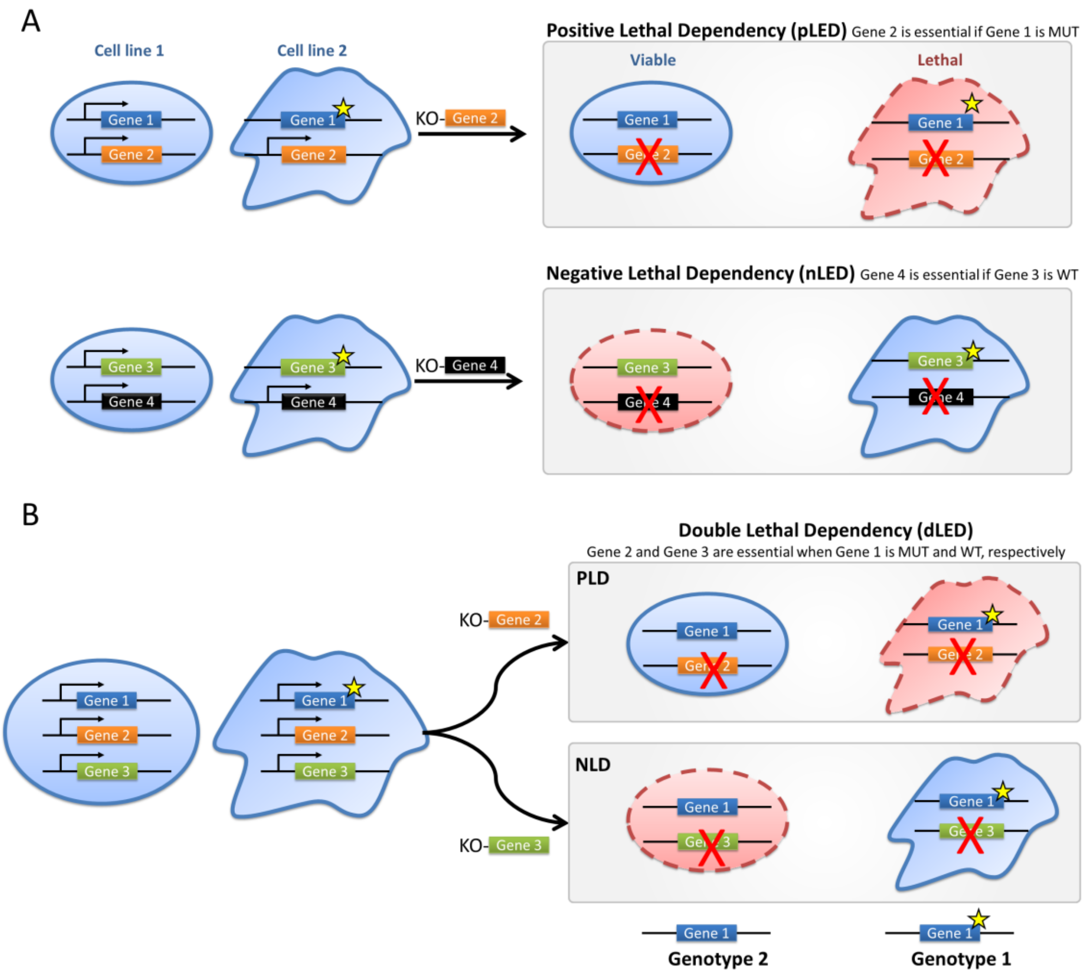

**Figure S1. Types of Lethal Dependencies.** Lethal dependencies that affect two genes: (A) in a Positive Lethal Dependency (pLED), a gene is essential for tumor survival when another gene is mutated (MUT). This is the traditional concept of synthetic lethality, in which a gene knockout (KO) causes cellular death only for another gene's mutant phenotype. (B) Conversely, in Negative Lethal Dependency (nLED), a gene is essential for tumor survival when another gene is not genetically altered (wild type-WT), here gene variant confers resistance to the inhibition. (C) A lethal dependency that affects three genes: Dual Lethal Dependency (dLED), an altered gene (Gene 1) confers, at the same time, sensitivity to the inhibition of one gene (Gene 2) and resistance to the inhibition of another gene (Gene 3). In this figure, the shape of the cells denotes different cell-types with different genomic characteristics. The color of the cells denote whether the cell survives to the knock down or not. The star shape in a gene denotes a genetic variant. The red crosses denote pharmacological inhibition.

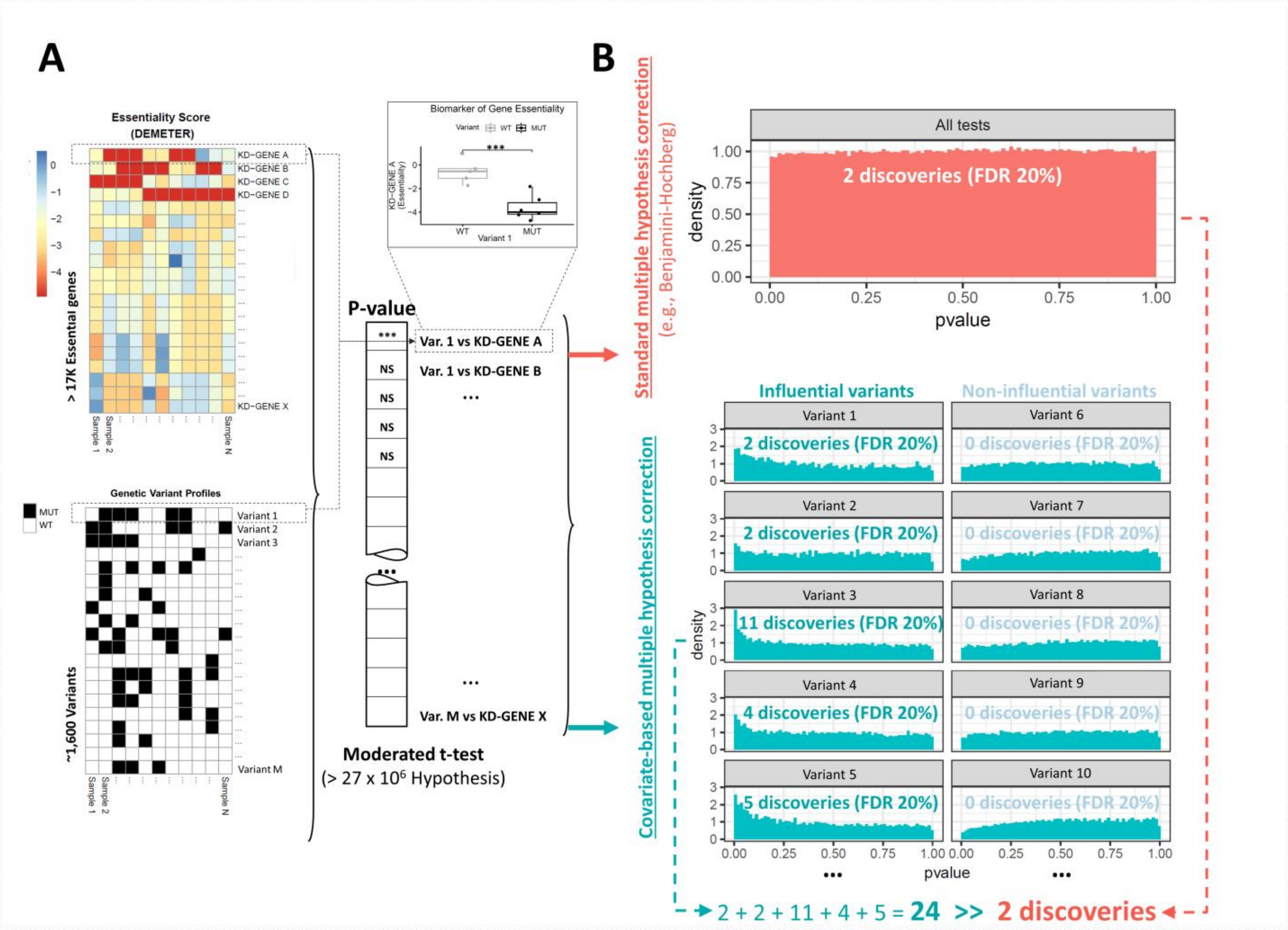

**Figure S2. Computational pipeline to find lethal dependencies.** (A) Scheme of data integration for N samples (cell lines). RNAi libraries data (gene essentiality; DEMETER score) and gene variant panels are represented as two heatmaps. Each pair of a knock-down gene (KD-gene) and a gene variant defines a p-value, which represents a lethal dependency. Boxplots represent the DEMETER score of a cell line when inhibiting one gene (X-axis) depending on the genetic alteration of another gene (Y-axis). Genes with a DEMETER score < -2 are considered essential for a cell line. (B) Scheme of the histogram of P-values using standard approaches (e.g., Storey-Tibshirani), in red; and using a covariate-based algorithm, in blue.

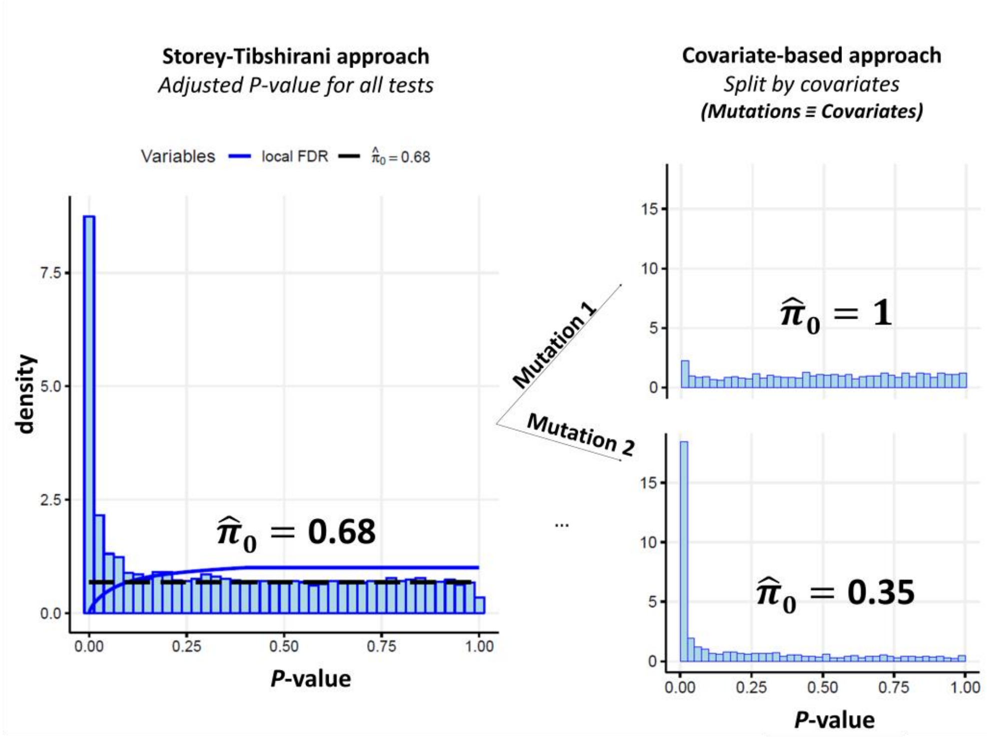

**Figure S3. Schematic representation of the covariate-based statistical approach in this context**. In this case, the use of genetic variants as covariates of a covariate-based problem allows the reduction of false positive rate, and consequently, the percentage of true null-hypothesis in a statistical test.

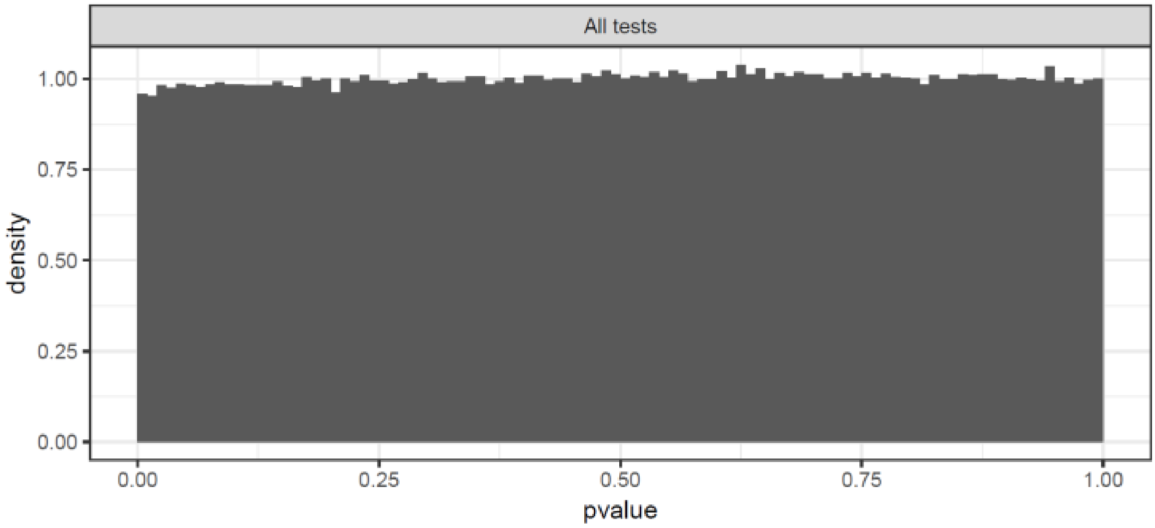

**Figure S4. Histogram of P-values of all lethal dependencies in acute myeloid leukemia.** Previous efforts to correct multiple testing in this problem consider a single set of tests (all gene aberrations and CRISPR-Cas9 knockouts) and apply a correction that control the FDR, such as Storey-Tibshirani (ST), as done in the Project Score. Interestingly, in this approach histogram of p-values shows flat-shaped histograms.

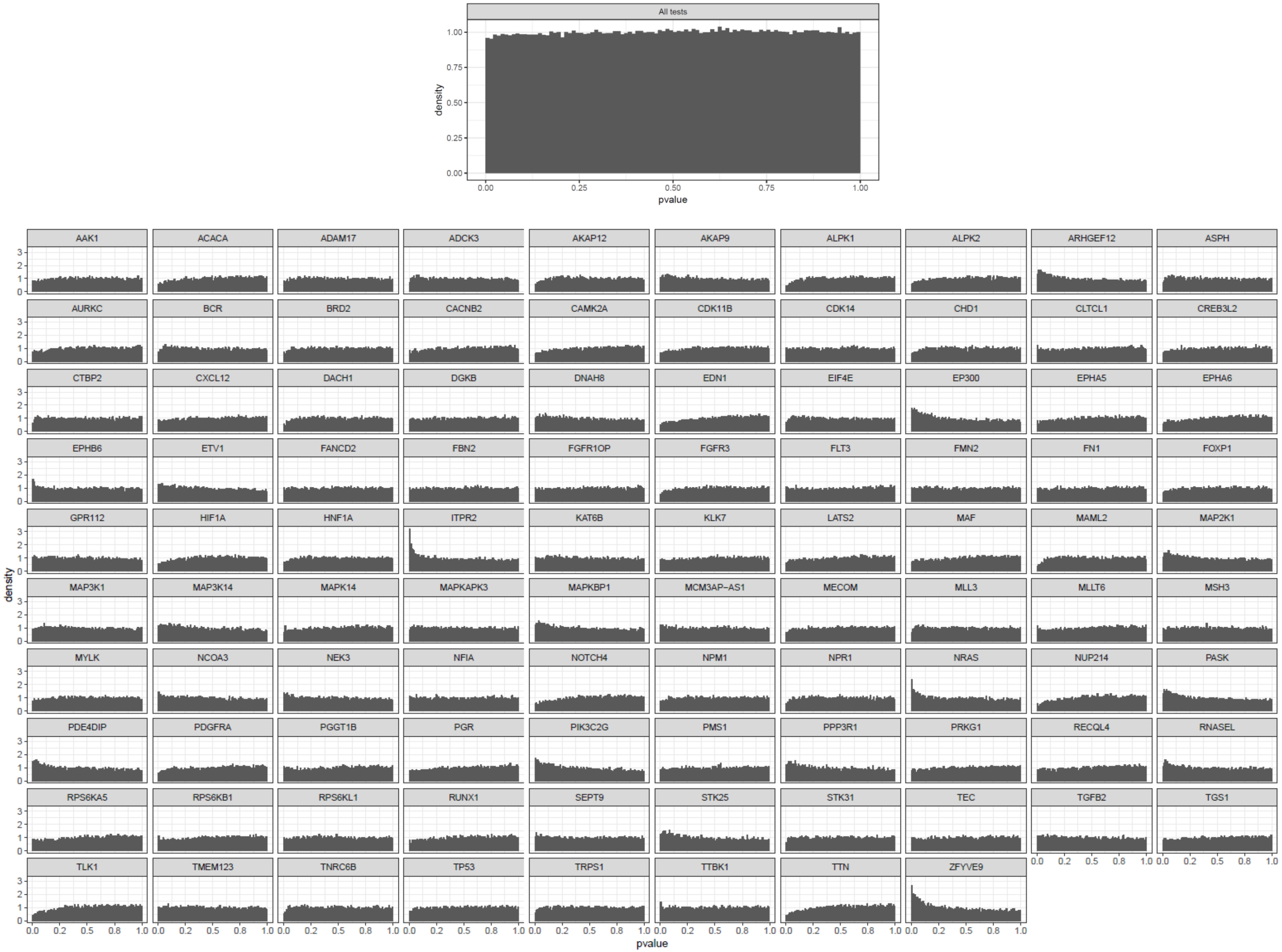

**Figure S5. Histogram of P-values of all lethal dependencies in acute myeloid leukemia vs p-values associated with each gene variant.**

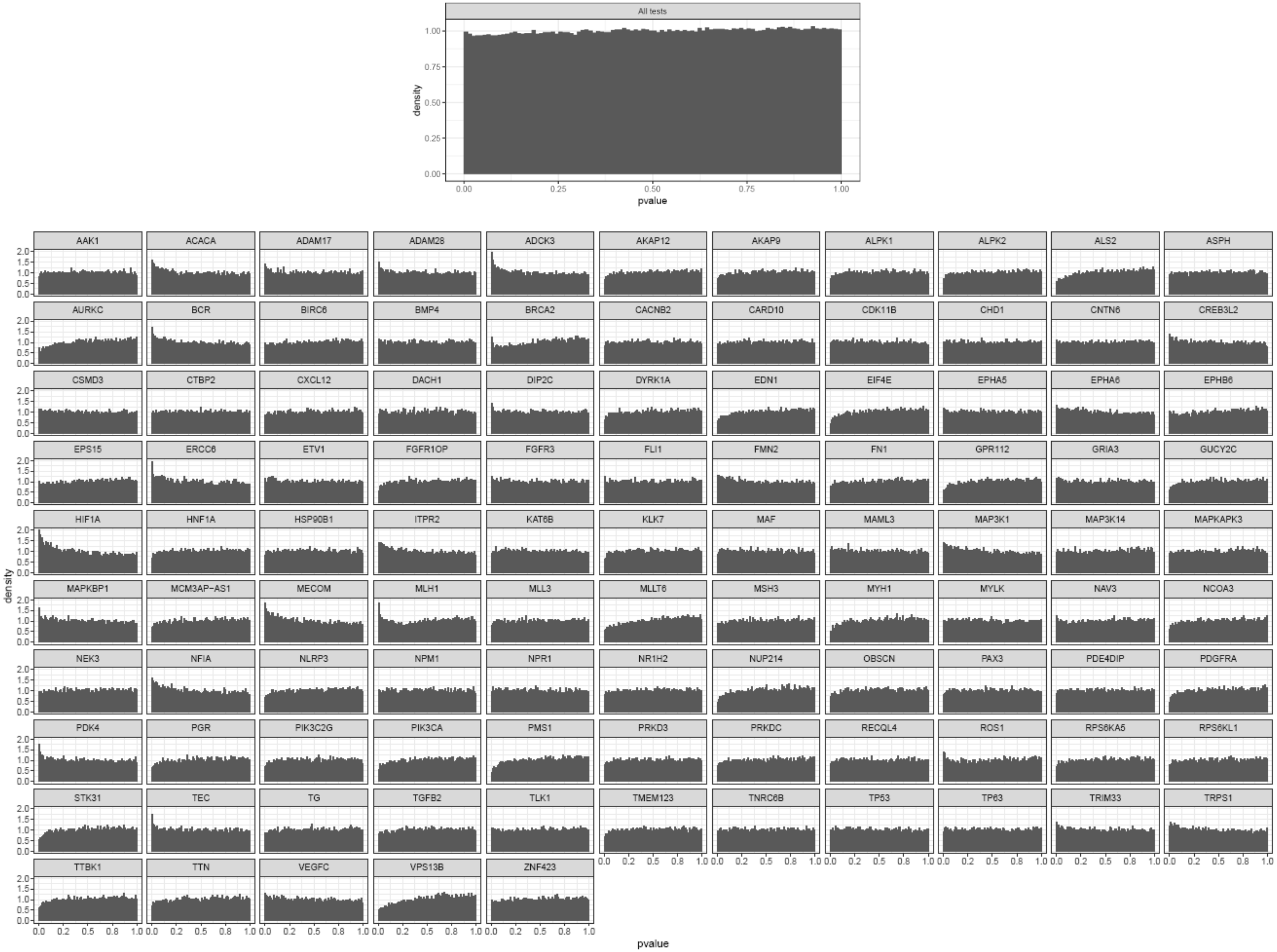

**Figure S6. Histogram of P-values oflethal dependencies in breast cancer vs p-values associated with each gene variant.**

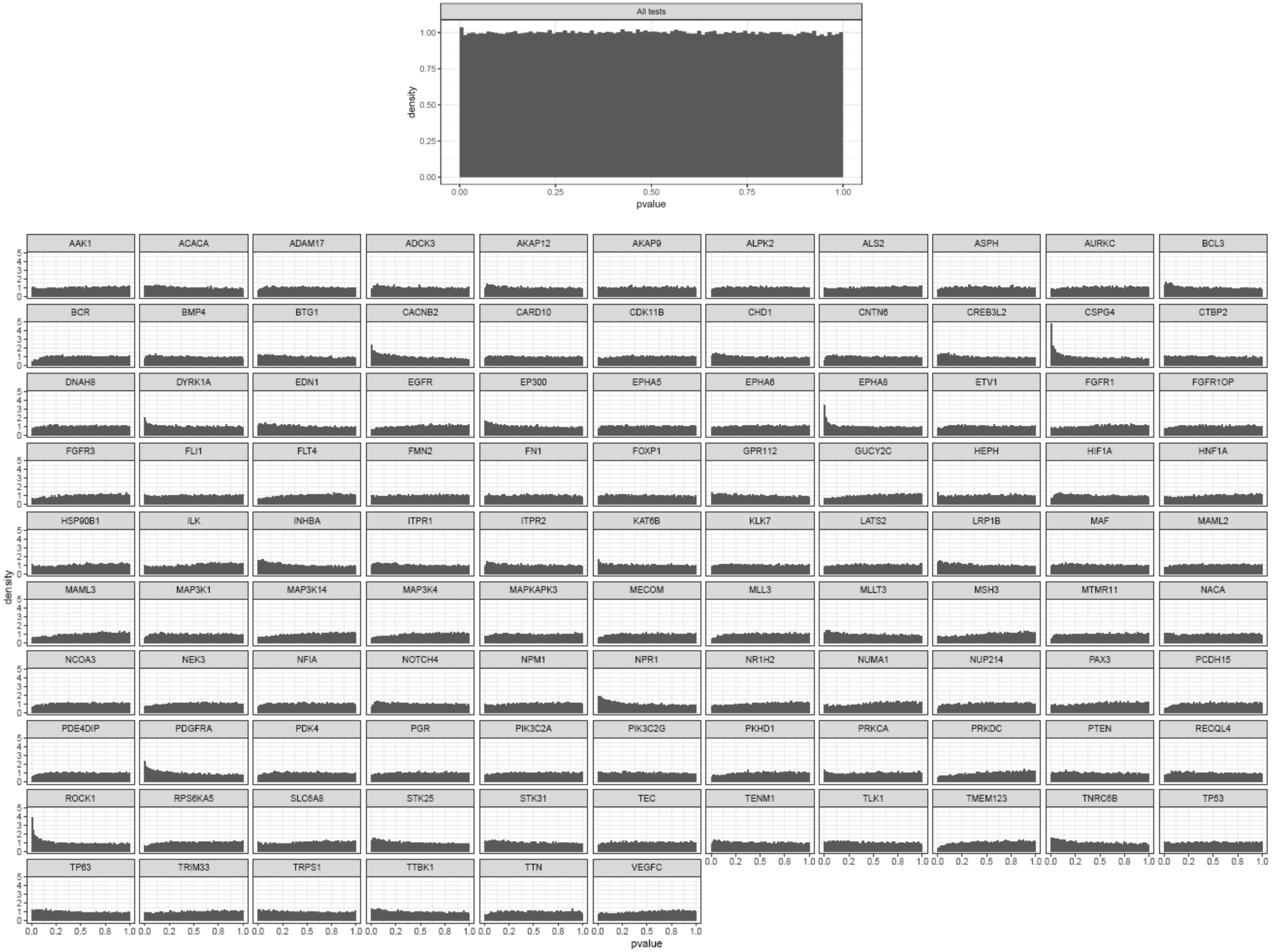

**Figure S7. Histogram of P-values of all lethal dependencies in central nervous system astrocytoma grade IV vs p-values associated with each gene variant.**

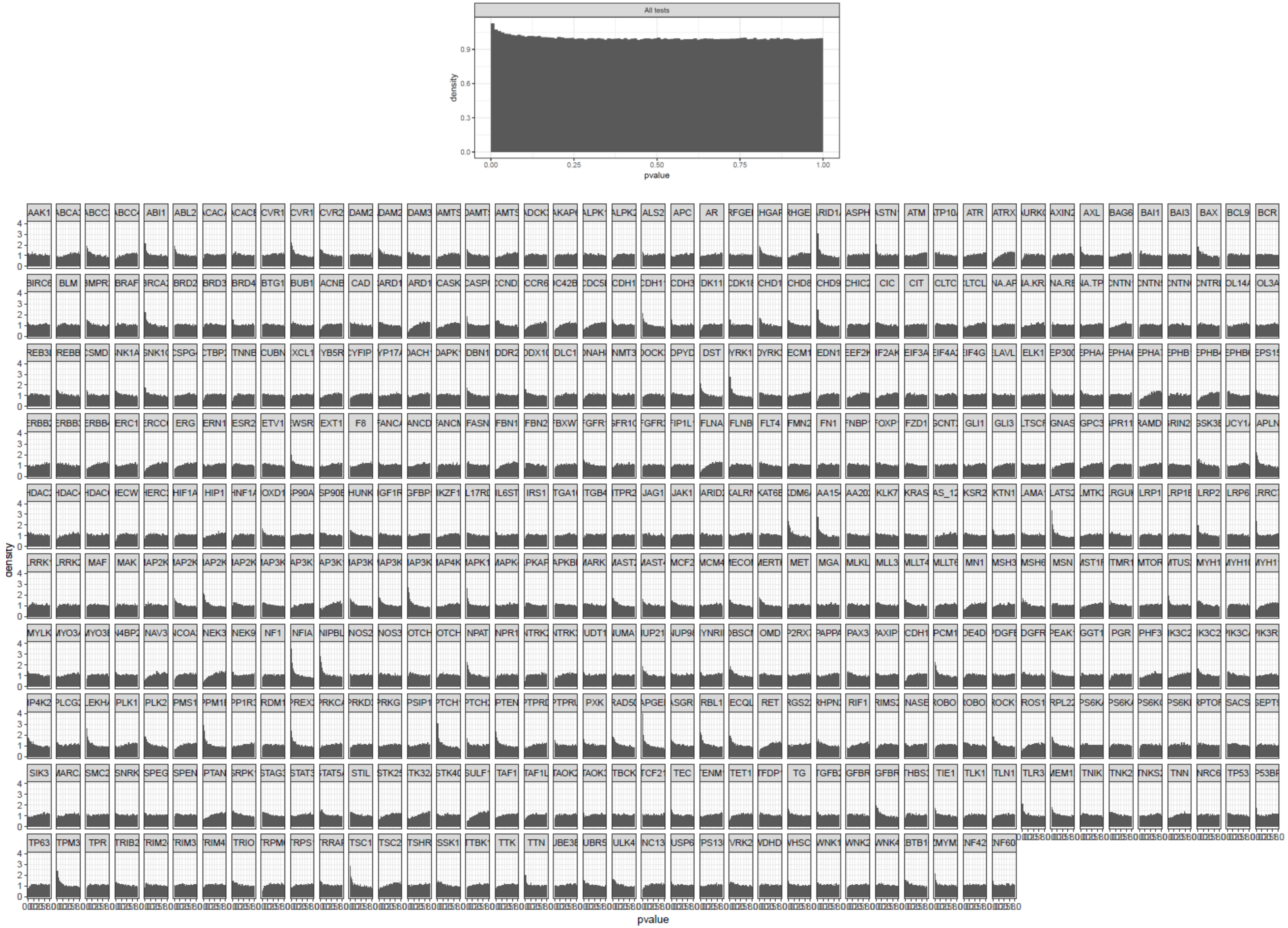

**Figure S8. Histogram of P-values of all lethal dependencies in colon adenocarcinoma vs p-values associated with each gene variant.**

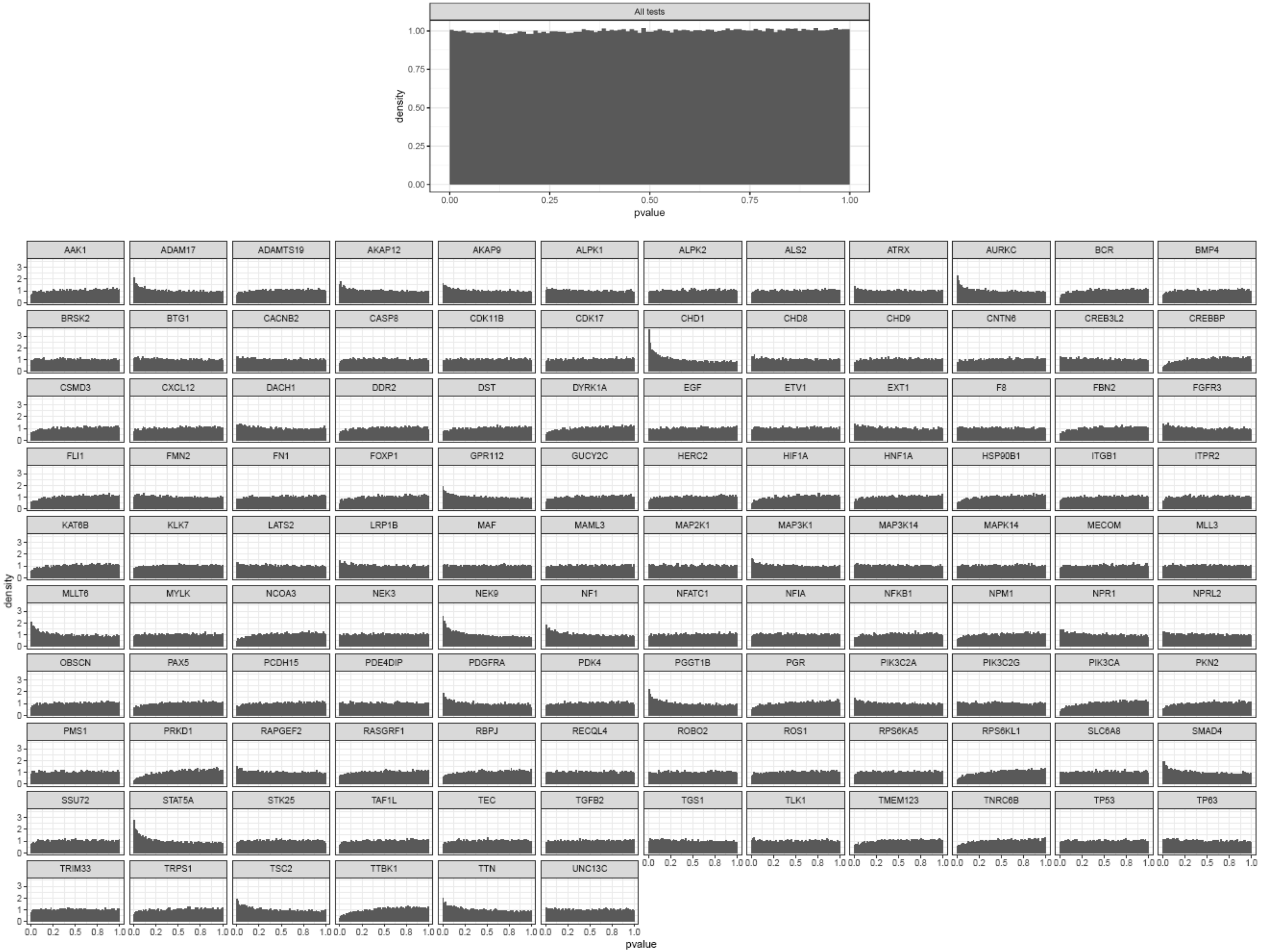

**Figure S9. Histogram of P-values of all lethal dependencies in upper aerodigestive tract squamous cell carcinoma vs p-values associated with each gene variant.**

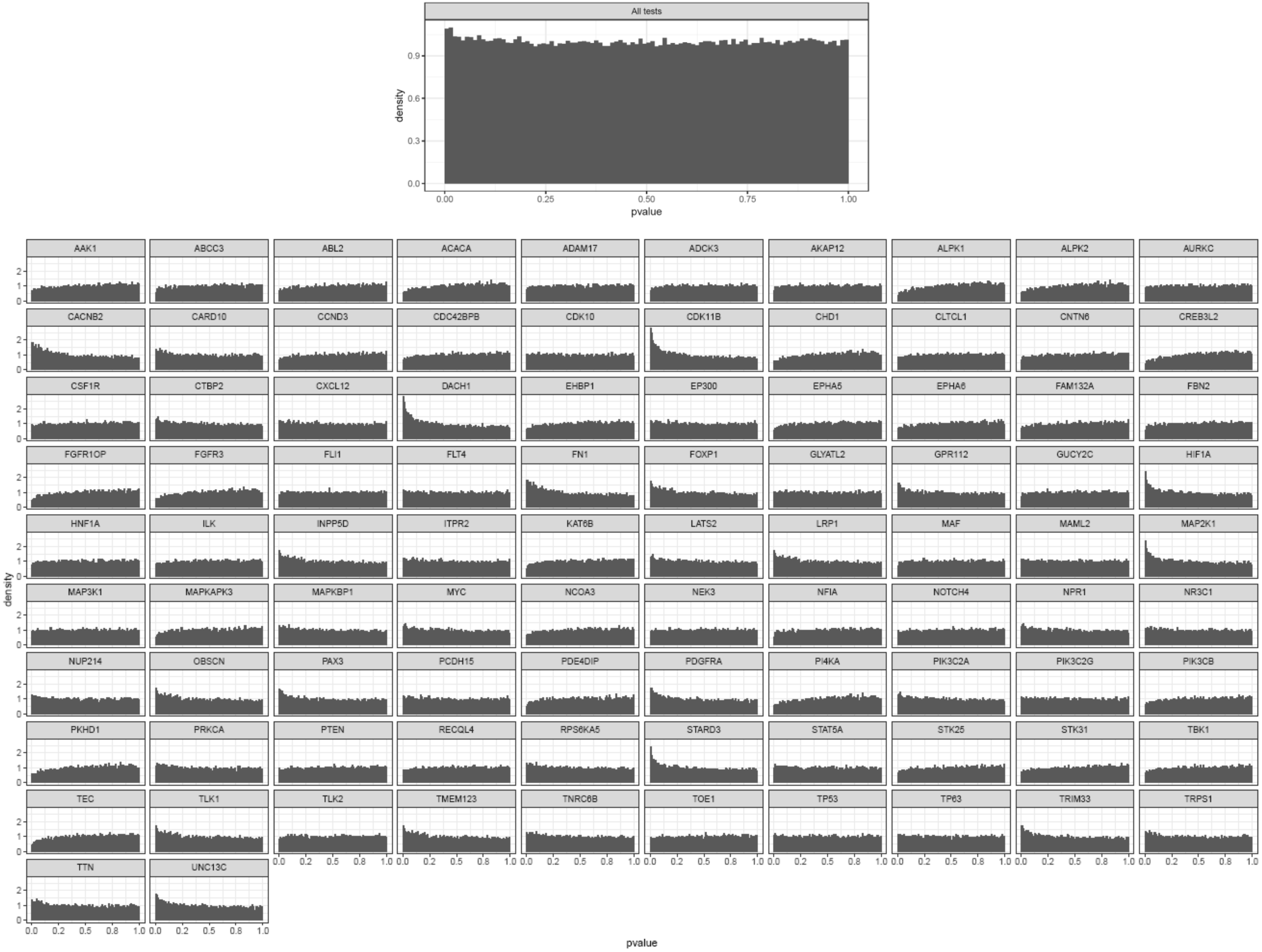

**Figure S10. Histogram of P-values of all lethal dependencies in diffuse large B-cell lymphoma vs p-values associated with each gene variant.**

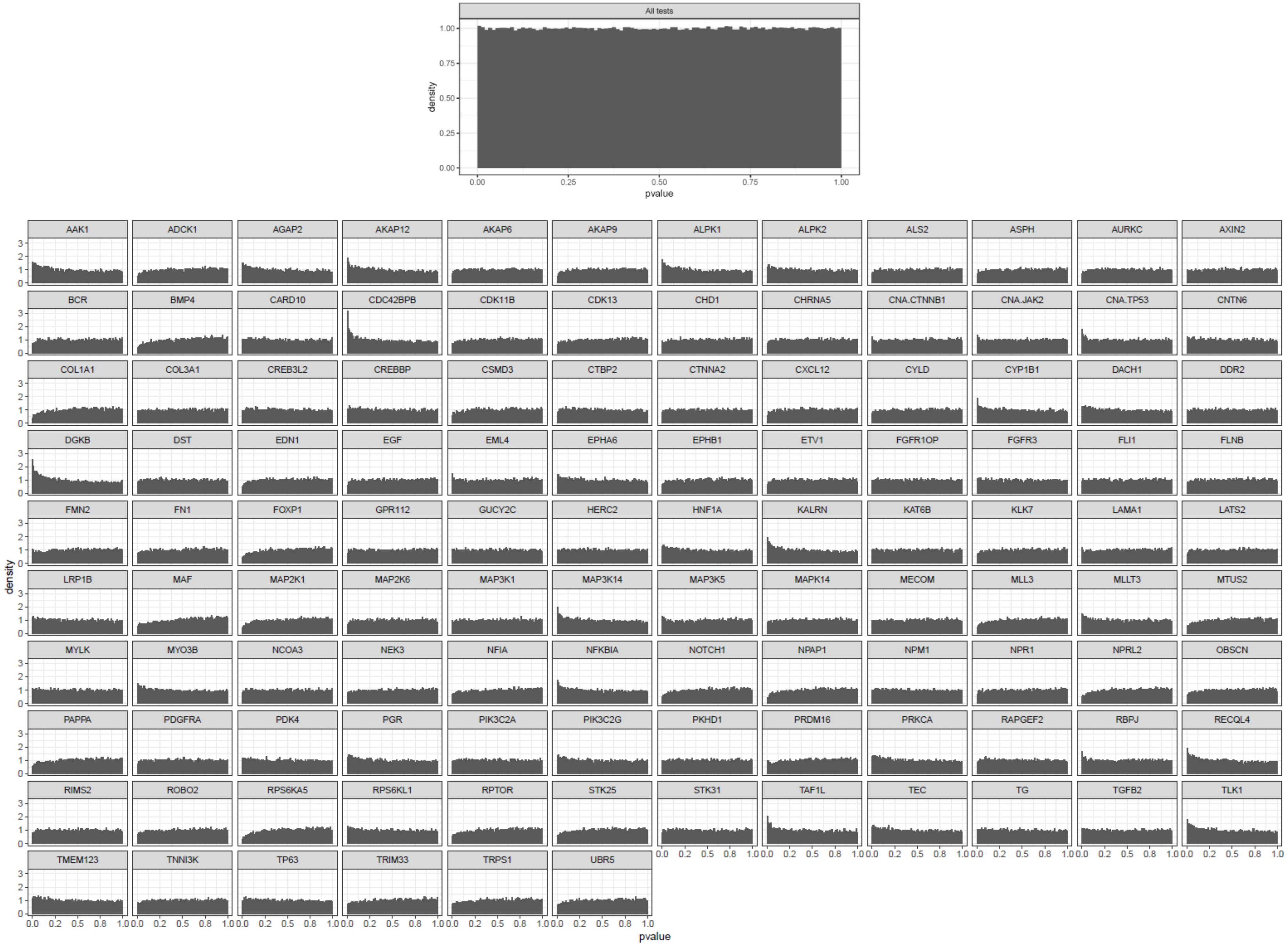

**Figure S11. Histogram of P-values of all lethal dependencies in esophagus squamous cell carcinoma vs p-values associated with each gene variant.**

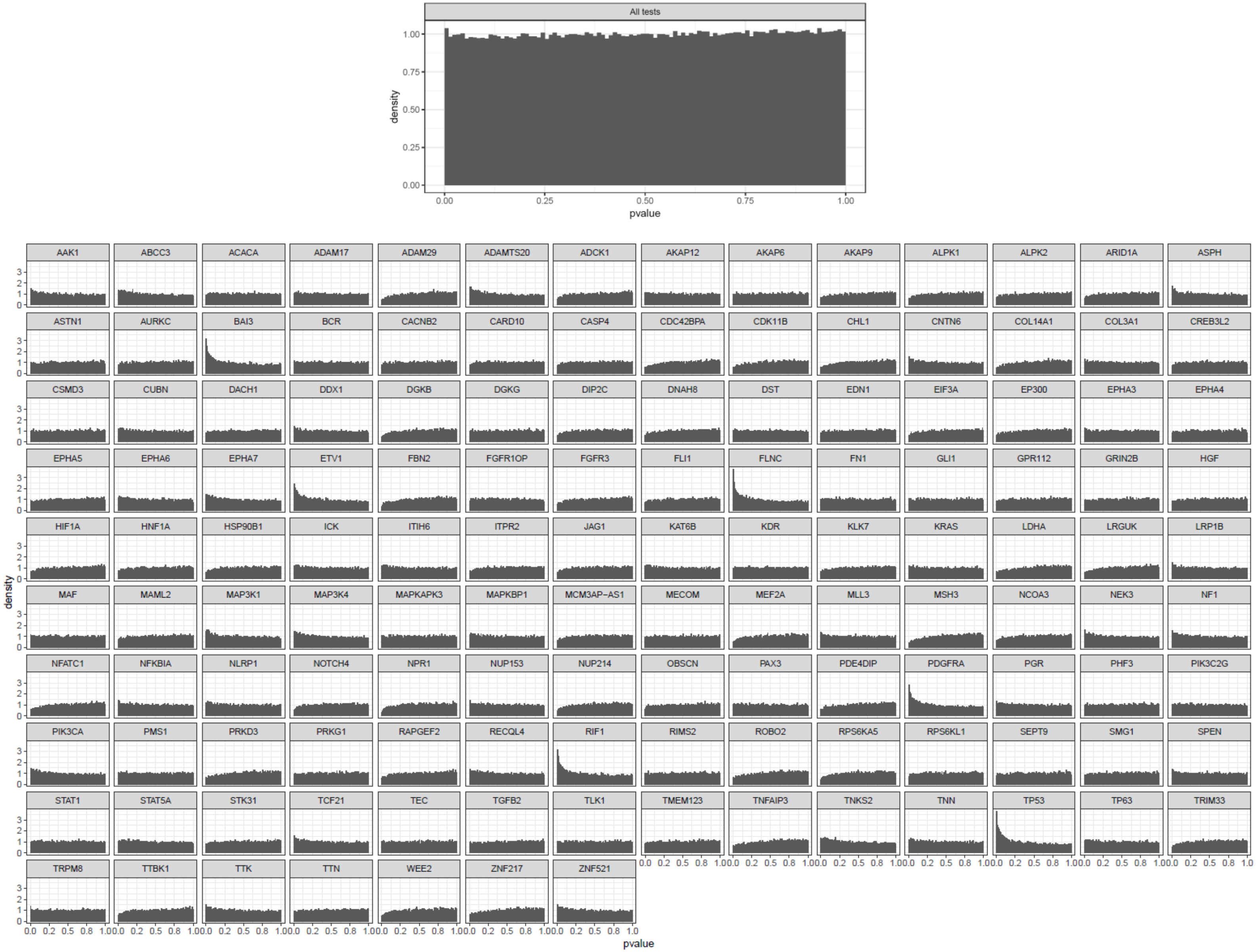

**Figure S12. Histogram of P-values of all lethal dependencies in lung large cell carcinoma vs p-values associated with each gene variant.**

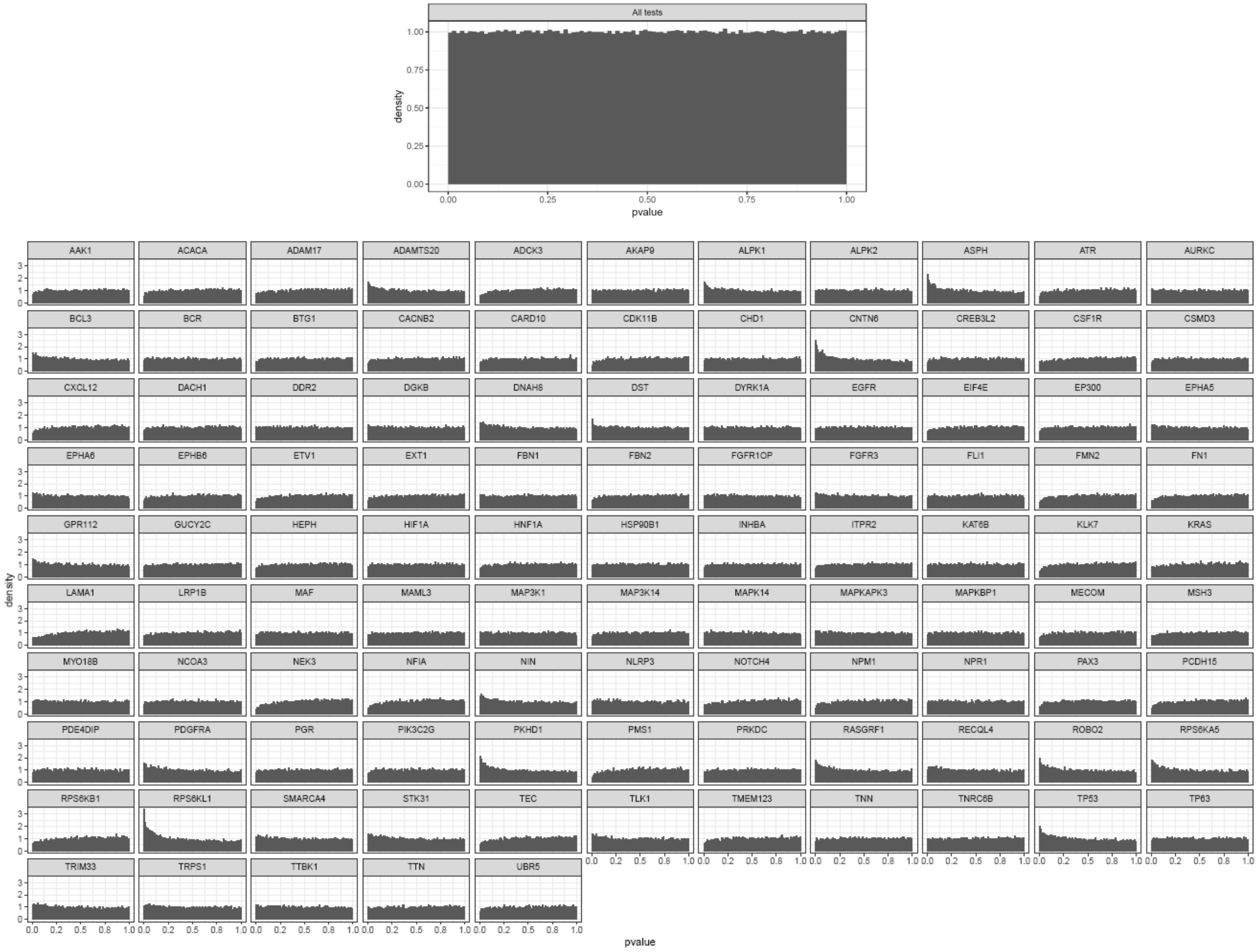

**Figure S13. Histogram of P-values of all lethal dependencies in lung adenocarcinoma vs p-values associated with each gene variant.**

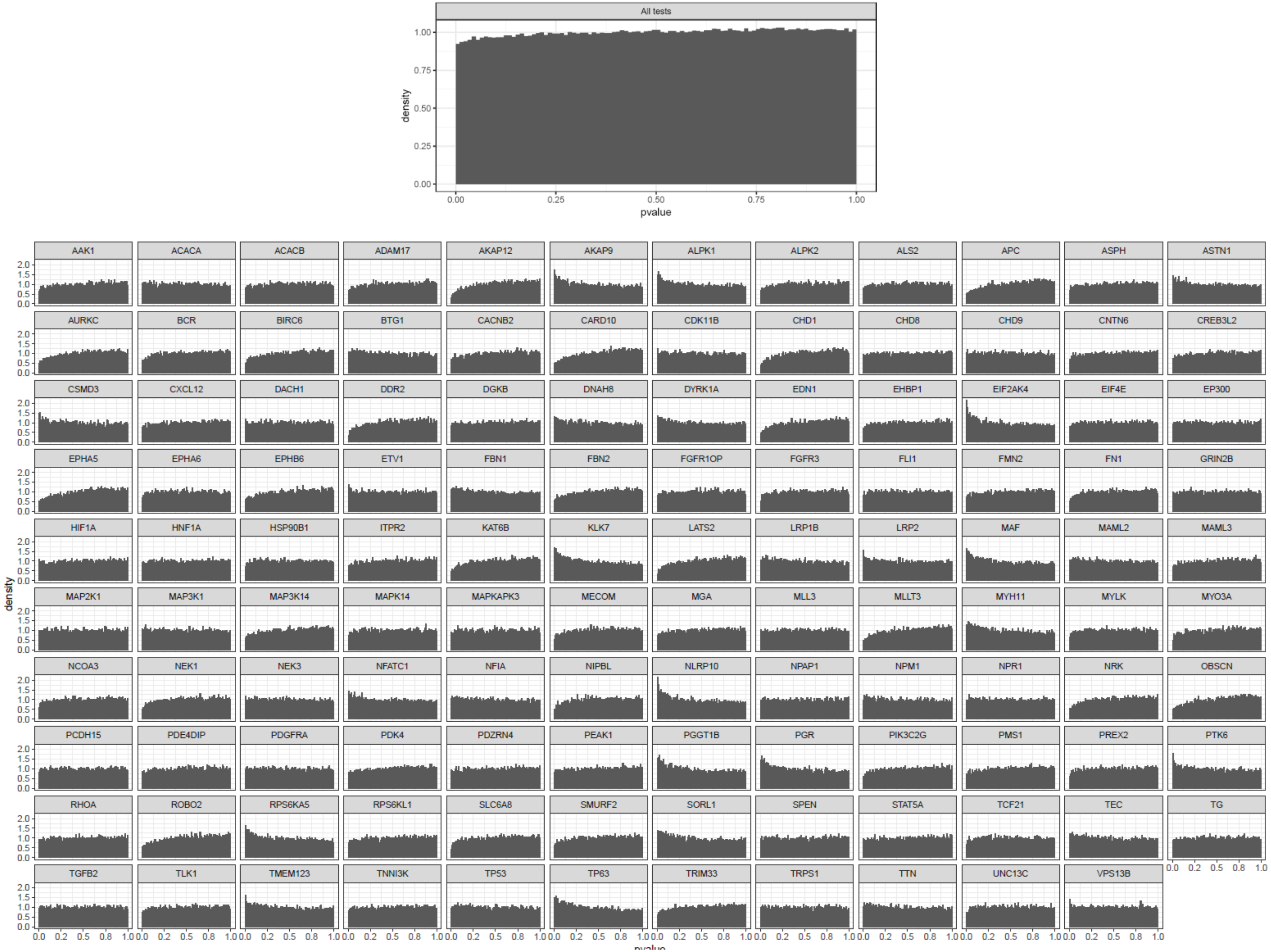

**Figure S14. Histogram of P-values of alllethal dependencies in lung squamous cell carcinoma vs p-values associated with each gene variant.**

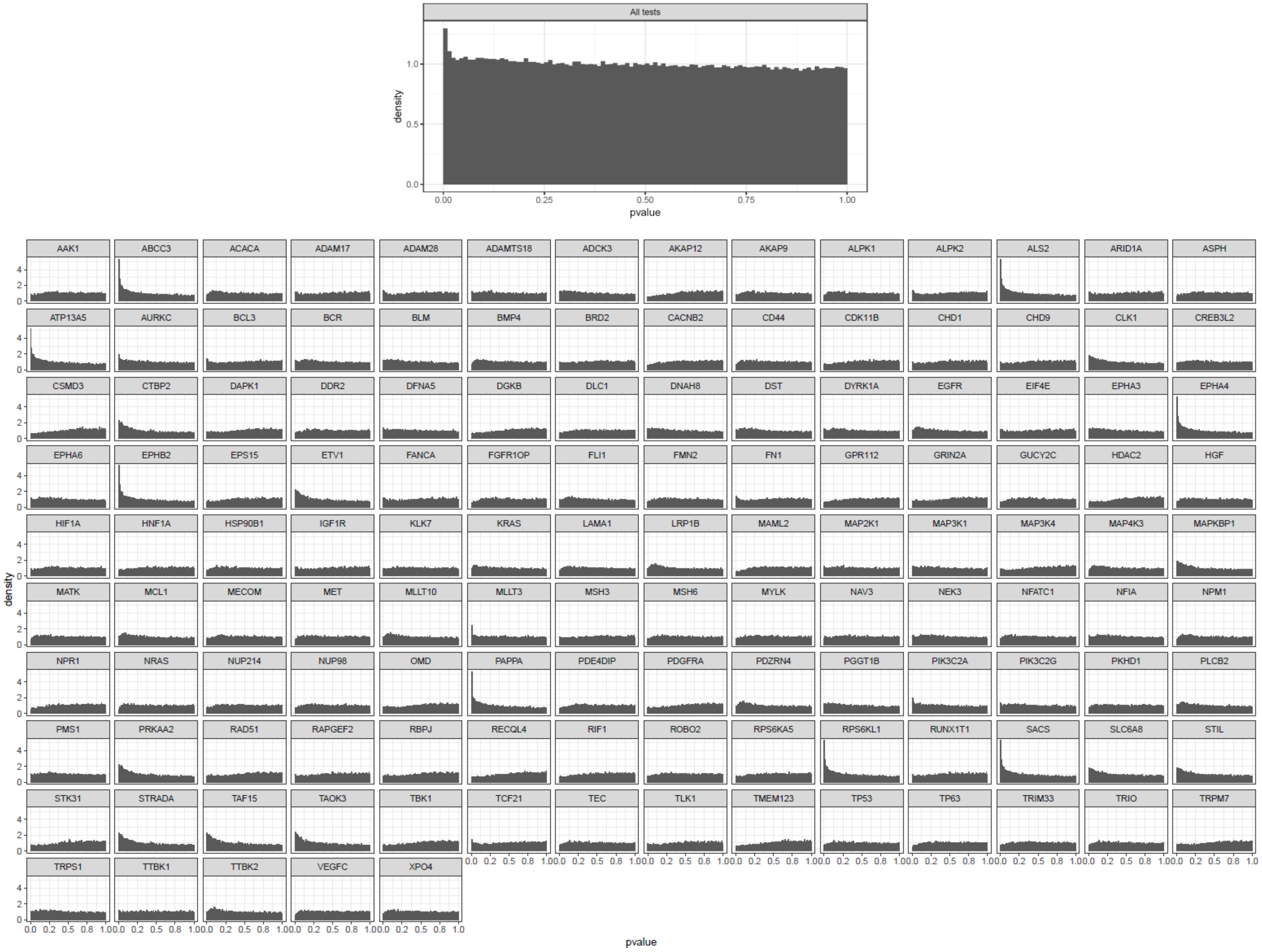

**Figure S15. Histogram of P-values of all lethal dependencies in multiple myeloma vs p-values associated with each gene variant.**

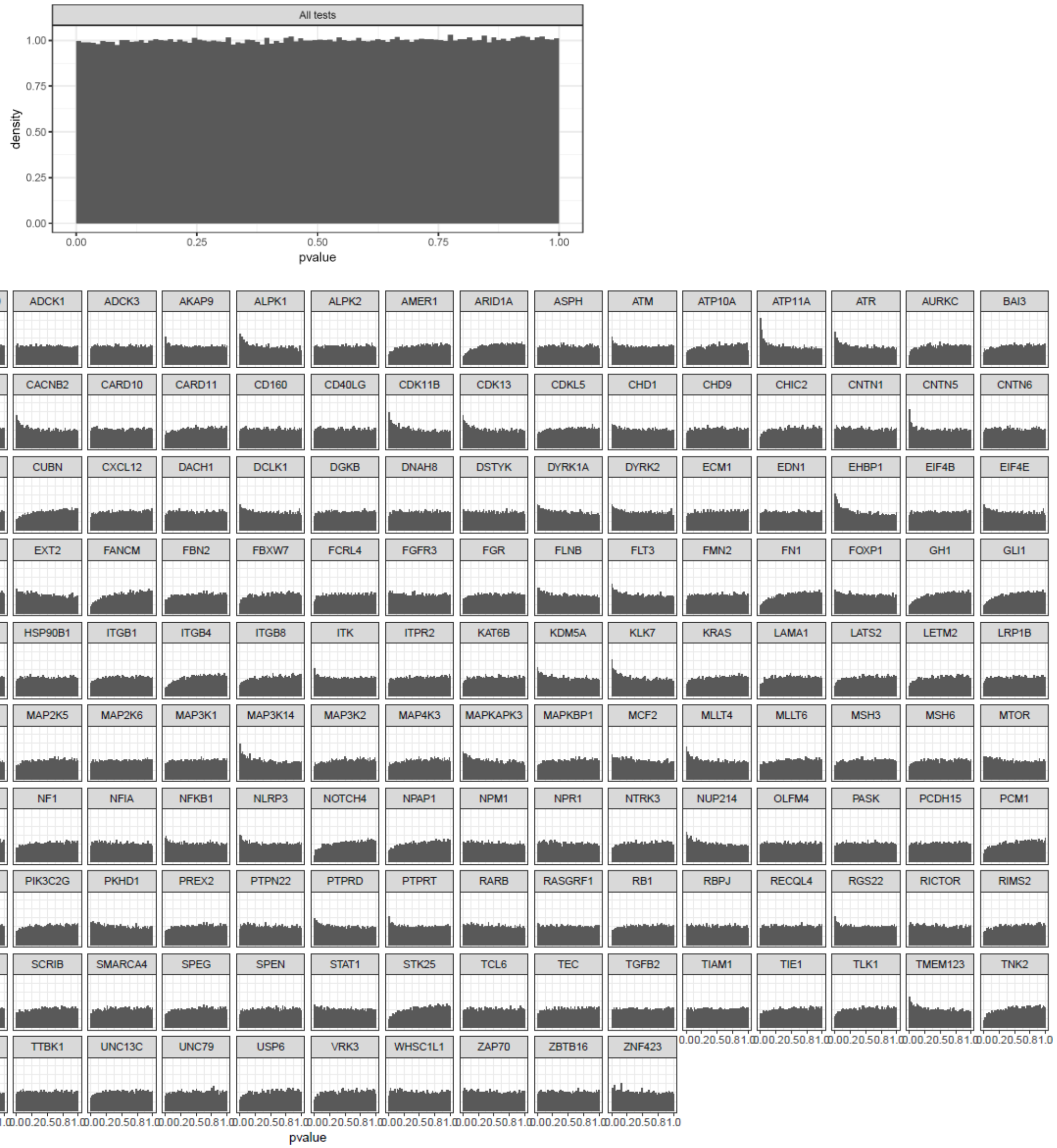

**Figure S16. Histogram of P-values of all lethal dependencies in non–small cell lung carcinoma vs p-values associated with each gene variant.**

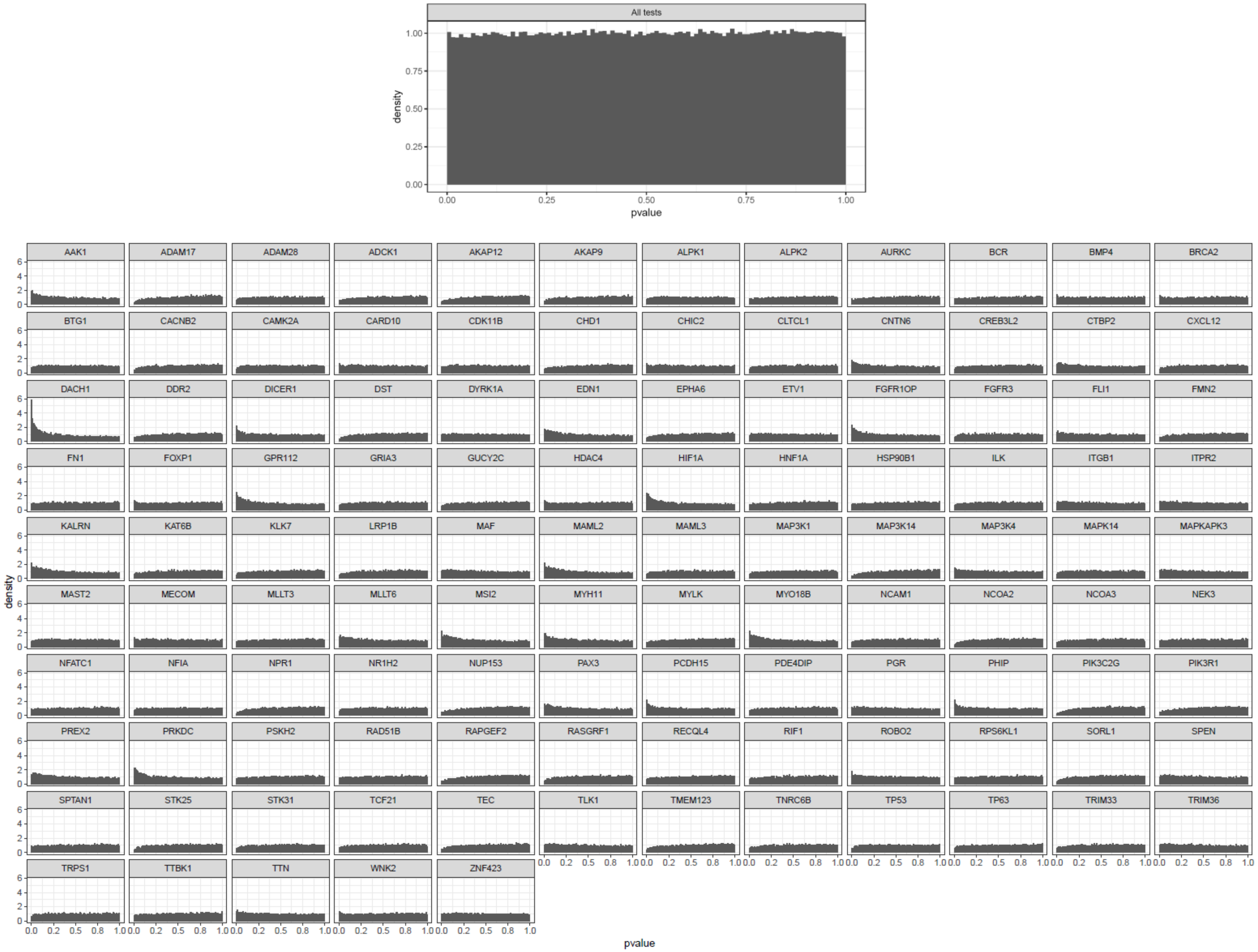

**Figure S17. Histogram of P-values of all lethal dependencies in osteosarcoma vs p-values associated with each gene variant.**

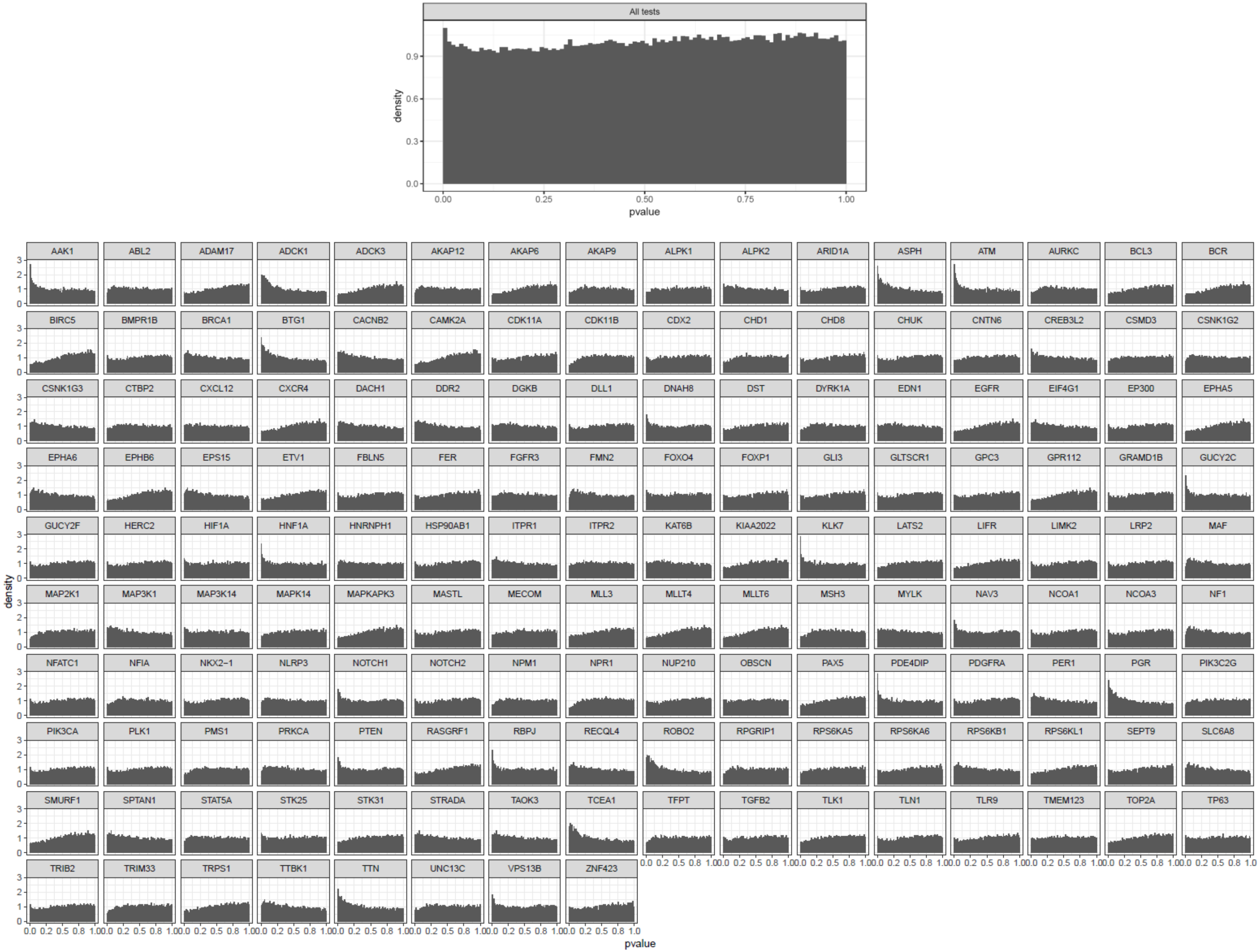

**Figure S18. Histogram of P-values of all lethal dependencies in ovary adenocarcinoma vs p-values associated with each gene variant.**

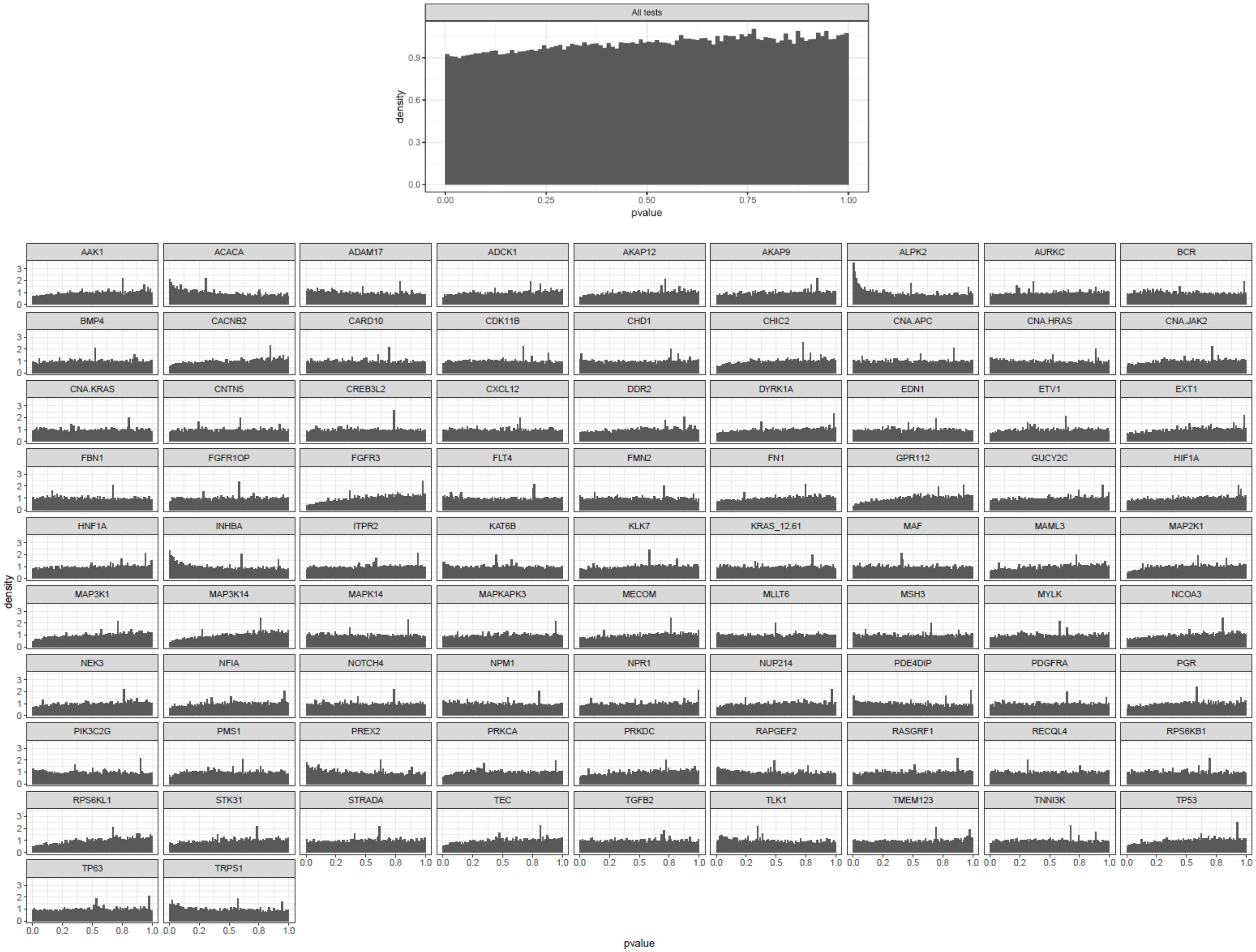

**Figure S19. Histogram of P-values of all lethal dependencies in pancreas ductal carcinoma vs p-values associated with each gene variant.**

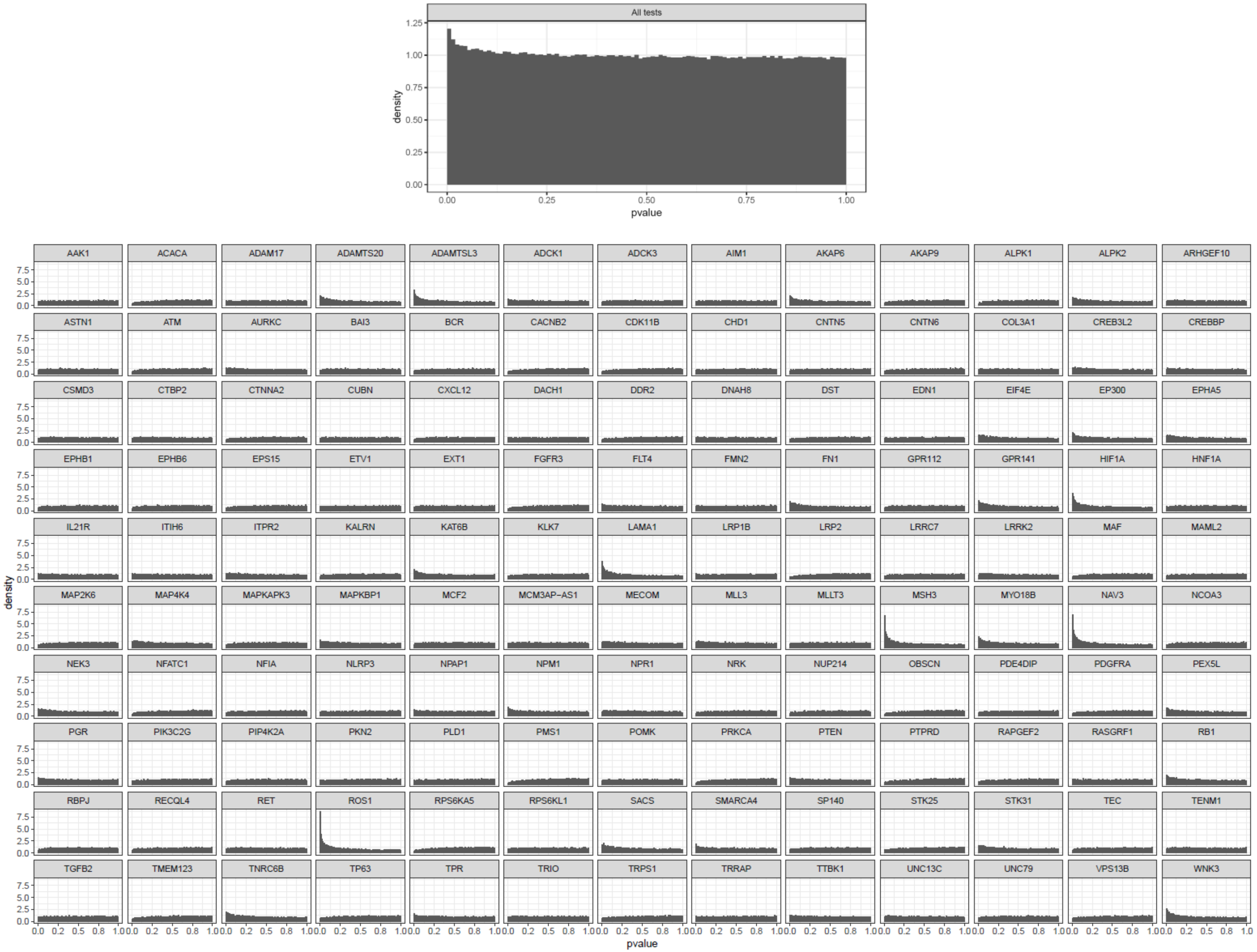

**Figure S20. Histogram of P-values of all lethal dependencies in small cell lung carcinoma vs p-values associated with each gene variant.**

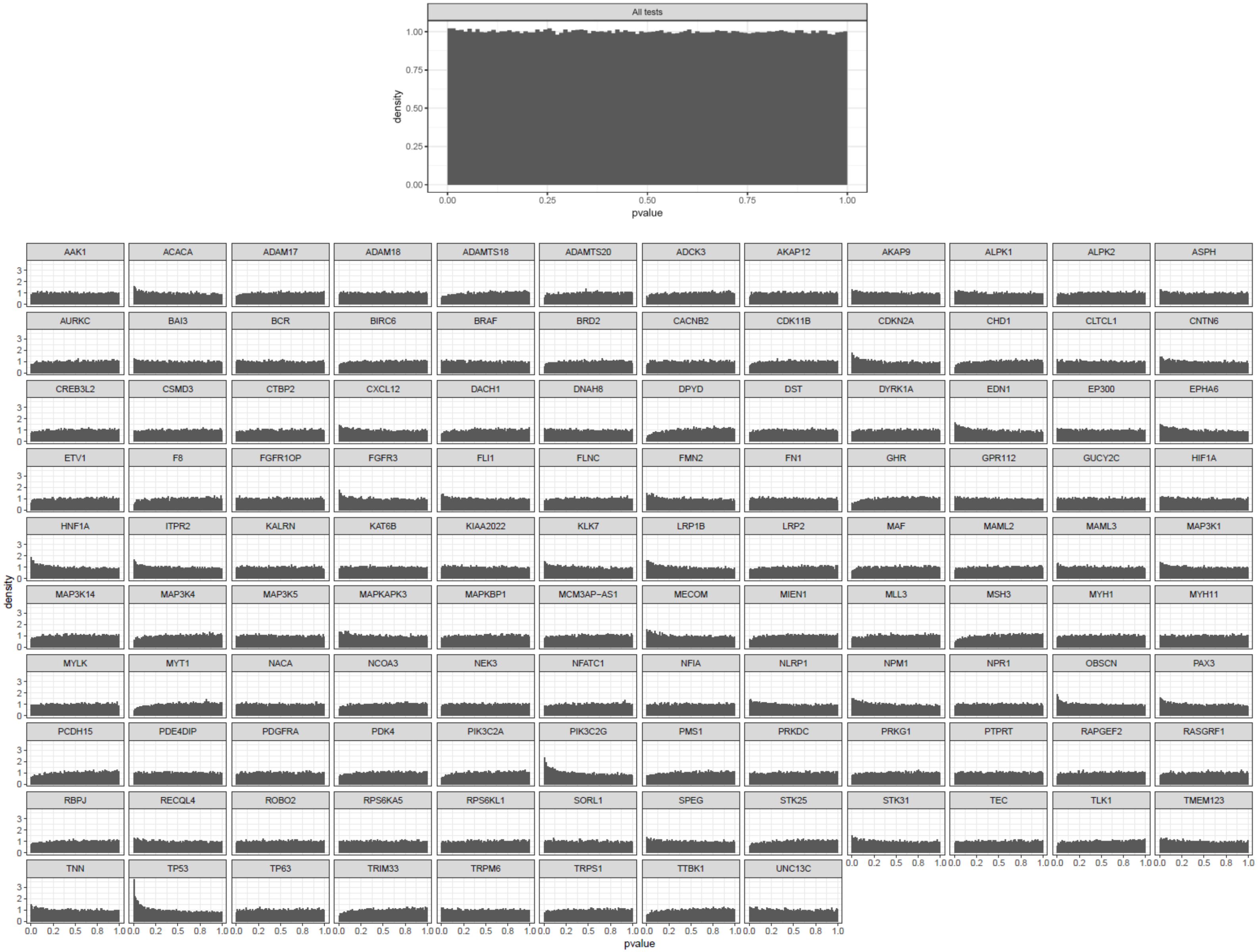

**Figure S21. Histogram of P-values of all lethal dependencies in skin carcinoma vs p-values associated with each gene variant.**

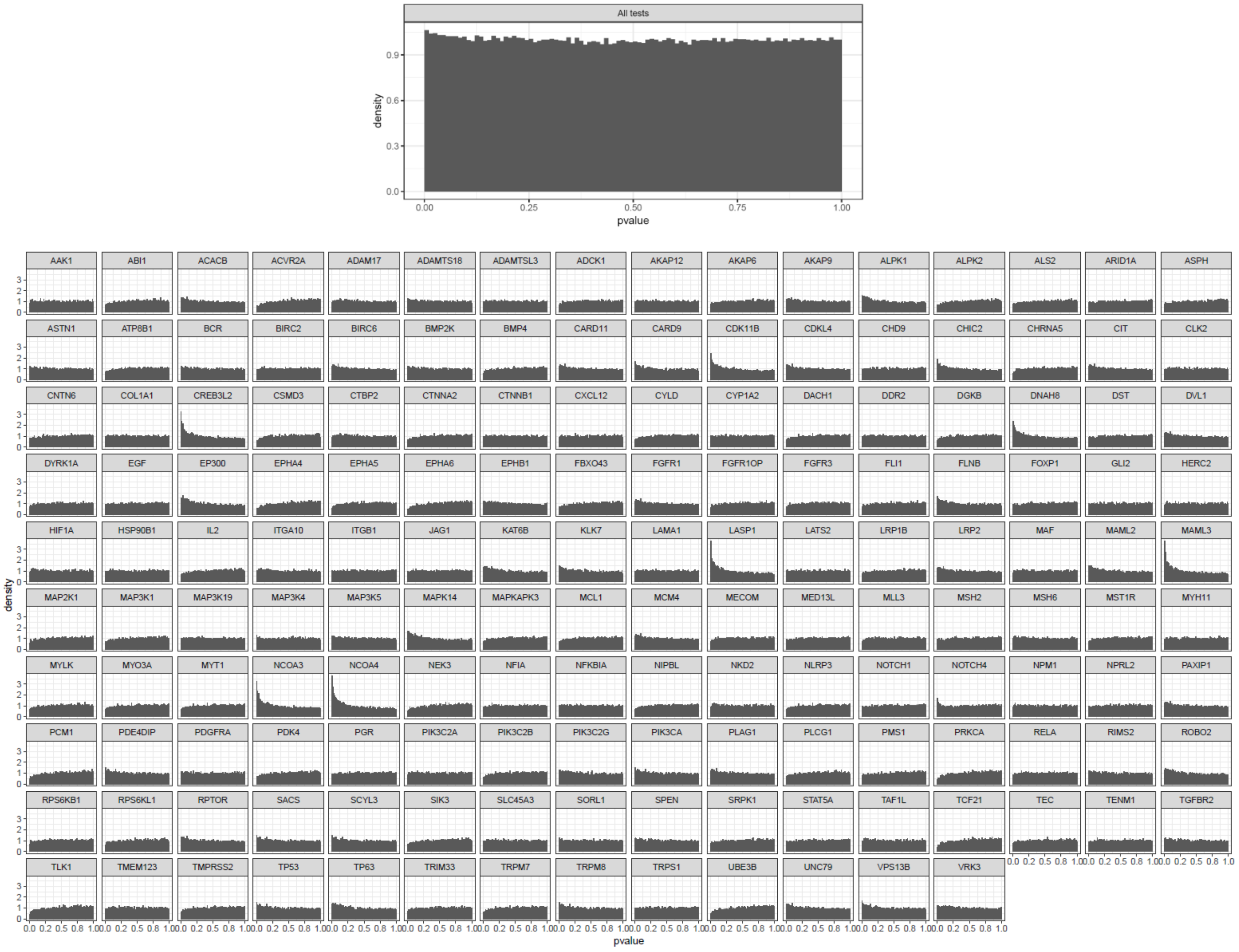

**Figure S22. Histogram of P-values of all lethal dependencies in stomach adenocarcinoma vs p-values associated with each gene variant.**

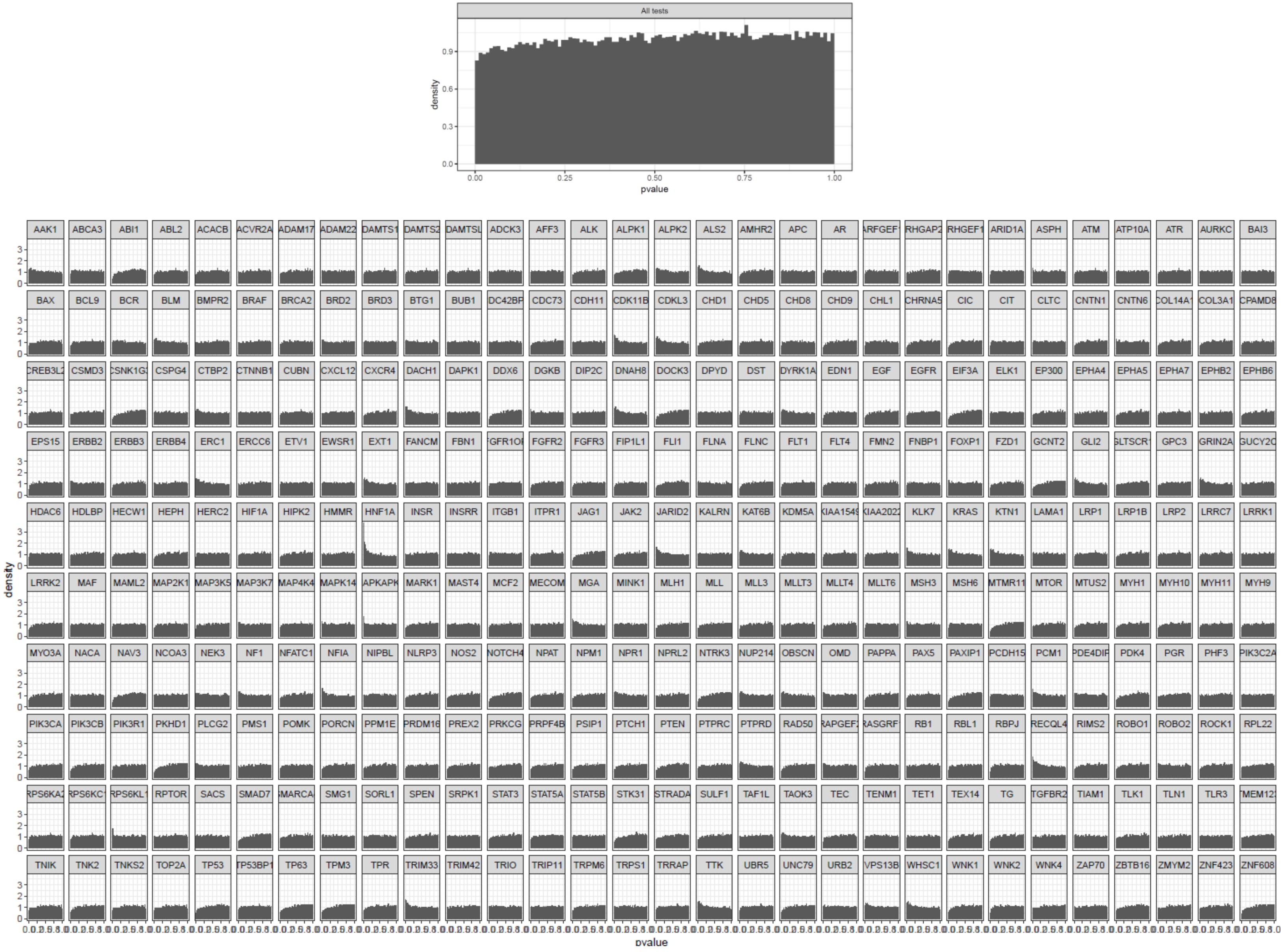

**Figure S23. Histogram of P-values of all lethal dependencies in uterine corpus endometrial carcinoma vs p-values associated with each gene variant.**

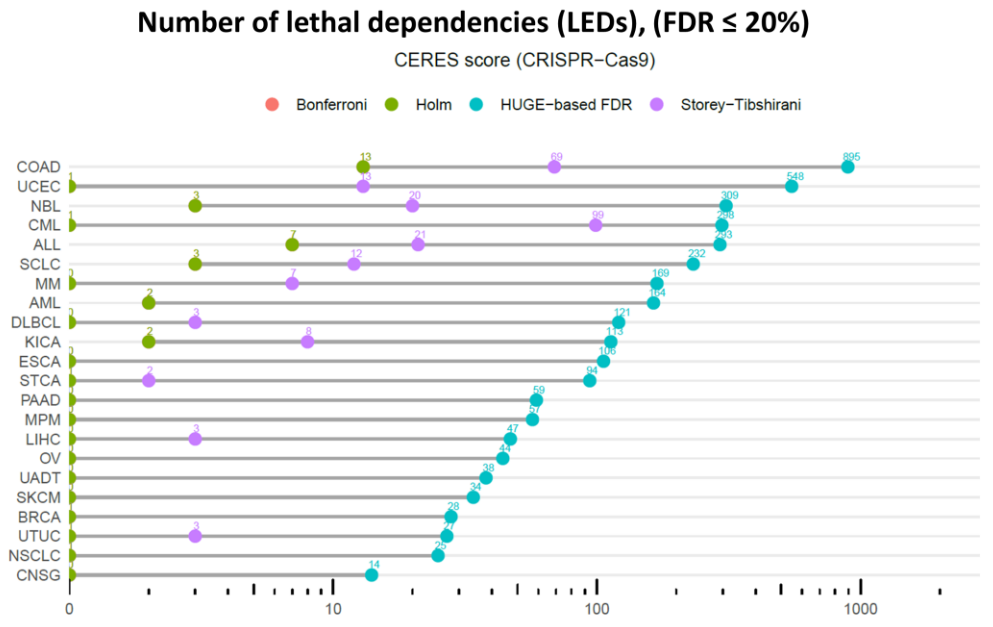

**Figure S24. The number of LEDs found (FDR ≤ 20%)** in 22 tumors of the CERES score (CRISPR-Cas9) using standard statistical pipelines (Storey-Tibshinary, Bonferroni, and Holm) and the HUGE-based algorithm. Bonferroni and Holm return the same number of hypotheses in all cases. LEGEND: ALL: acute lymphoblastic leukemia; AML: acute myeloid leukemia; BRCA: breast ductal carcinoma; CNSA-IV: central nervous system astrocytoma grade IV; COAD: colon adenocarcinoma; CUADT: upper aero-digestive tract squamous cell carcinoma; DLBCL: diffuse large B-cell lymphoma; ESCA: esophagus squamous cell carcinoma; KIRC: kidney renal clear cell carcinoma; LCC: lung large cell carcinoma; LUAD: lung adenocarcinoma; LUSC: lung squamous cell carcinoma; MM: multiple myeloma; NSCLC: non–small cell lung carcinoma; OS: osteosarcoma; OVAD: ovary adenocarcinoma; PDAC: pancreas ductal carcinoma; SCLC: small cell lung carcinoma; SKCM: skin carcinoma; UCEC: endometrium adenocarcinoma.

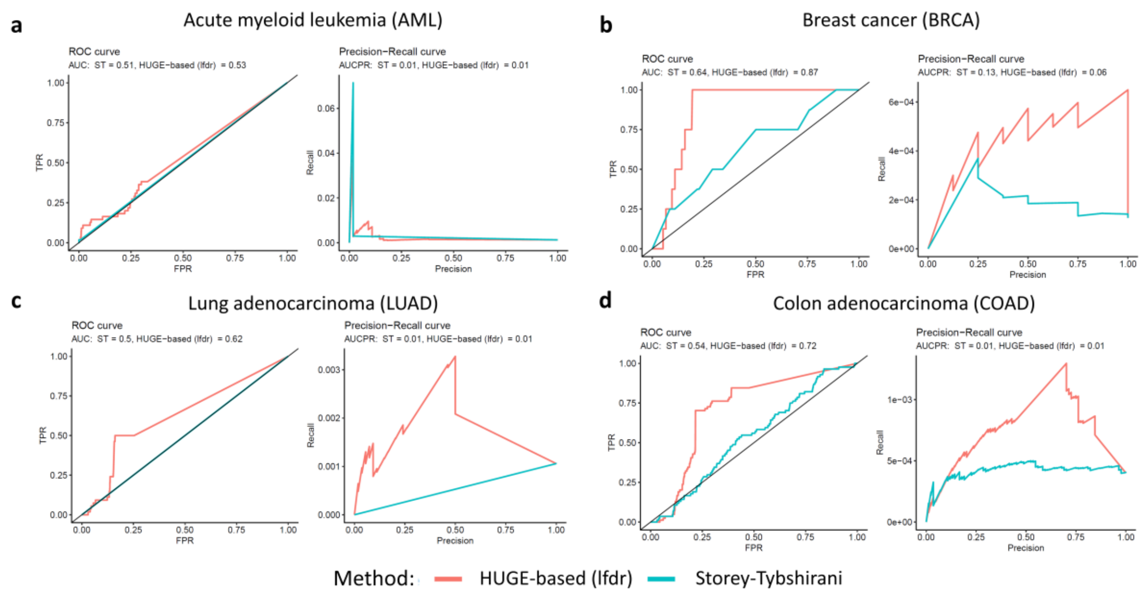

**Figure S25. ROC and precision-recall curves of four tumor types.** True positives are extracted from the somatic genomic variants in the knowledgebase of the Variant Interpretation for Cancer Consortium (VICC) for four different cancers. Each of the panels show a) AML, b) BRCA, c) LUAD and d) COAD. We selected associations indicated for each tumor type that are within the three highest levels of confidence (Level A: Evidence from professional guidelines or FDA-approved therapies relating to a biomarker and disease; Level B: Evidence from clinical trials or other wellpowered studies in clinical populations, with expert consensus; and Level C: Evidence for therapeutic predictive markers from case studies, or other biomarkers from several small studies, or evidence for biomarker therapeutic predictions for established drugs for different indications). Both the ROC and the Precision Recall curves have larger area for HUGE than fot the ST method.

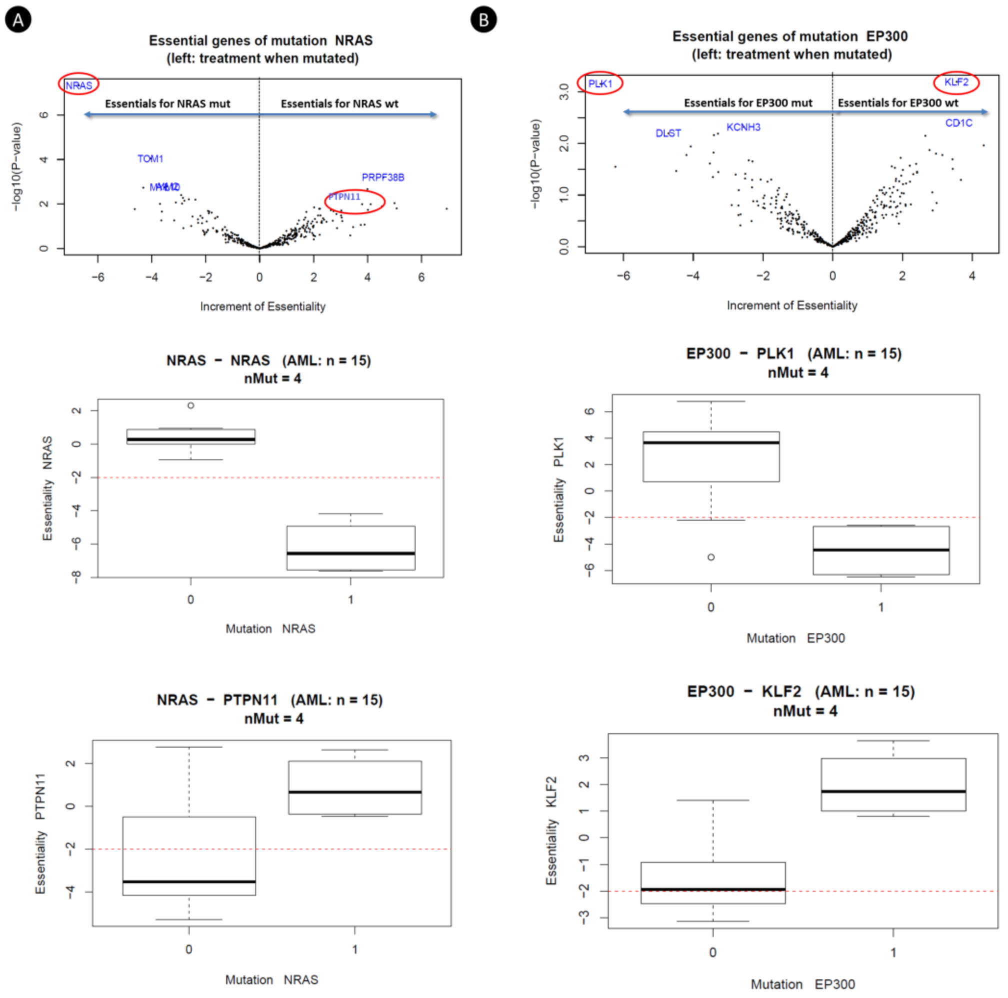

**Figure S26.** Volcano plot of Synthetic lethal genes related to *NRAS*-mutated (A) and *EP300*-mutated (B) phenotypes. Increment of Essentiality and -log10 (P-value) are shown y x-axis and y-axis respectively. Boxplots of AML P-values of same genes' synthetic lethal: *NRAS, PTPN11, PLK1* and *KLF2.*

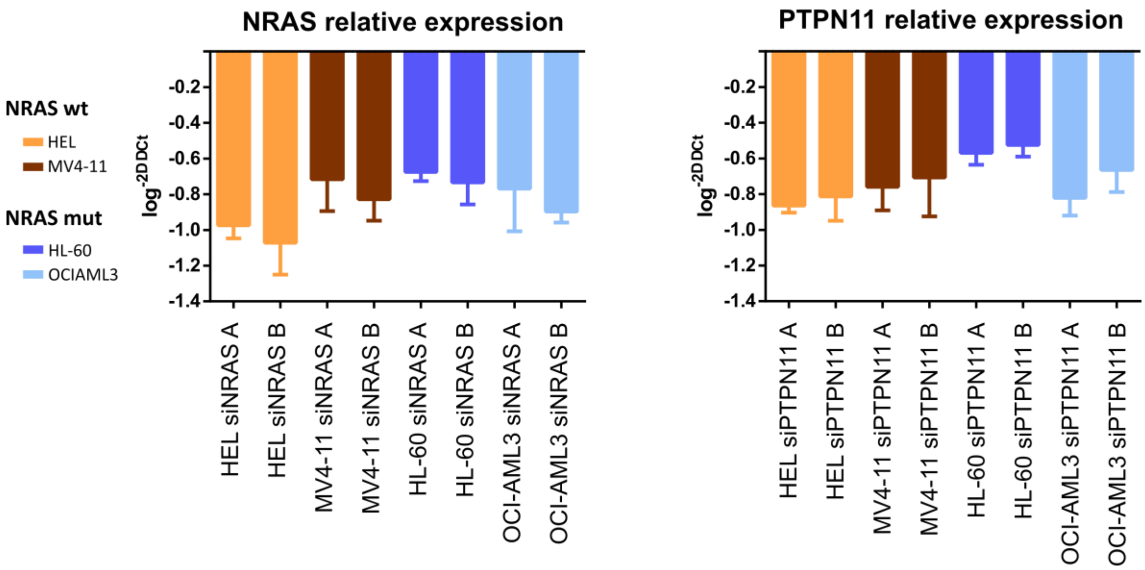

**Figure S27. mRNA expression of *NRAS* and *PTPN11* genes after nucleofection with the specific siRNAs.** Data are referred to *GUSB* gene and an experimental group nucleofected with negative control siRNA.

### SUPPLEMENTAL METHODS

#### Section 1. Cell culture

The AML cell lines HL-60, HEL, MV4-11 and OCI-AML3 were maintained in culture in RPMI-1640 medium supplemented with 10% fetal bovine serum (Gibco, Grand Island, NY), penicillin/streptomycin (BioWhitaker, Walkersvill, MD) at 37 °C in a humid atmosphere containing 5% CO2. All cell lines were tested for mycoplasma (MycoAlert Sample Kit, Cambrex) and were authenticated by performing a short tandem repeat allele profile.

#### Section 2. Cell transfection

Cells were passaged 24 hours before nucleofection, and cells for nucleofection were in their logarithmic growth phase. The transfection of siRNAs was done with the Nucleofector II device (Amaxa GmbH, Köln, Germany) following the Amaxa guidelines. Briefly, $1 \times 10^6$ of HL-60, HEL, MV4-11 and OCI-AML3 cells were resuspended in 100 µL of supplemented culture medium or solution V in the case of HL-60 cells, with 75nM of NRAS or PTPN11 siRNAs or Silencer Select Negative Control-1 siRNA (Ambion, Austin, TX) and nucleofected with the Amaxa nucleofector apparatus using programs A030 (HEL, MV4-11 and OCI-AML3) or T019 (HL-60). We used two different siRNAs against *NRAS* target (siNRAS A: GAACCACUUUGUAGAUGAA; siNRAS B: AAGGACAGTTGATACAAAA) and *PTPN11* (siPTPN11 A: AGAUGUCAUUGAGCUUAAA; siPTPN11 B: GAAAGAAGCAGAGAAAUUA) to demonstrate that the results obtained with siRNA nucleofection are not due to a combination of inconsistent silencing and sequence specific off-target effects. Silencer Select Negative Control-1 siRNA was used to demonstrate that the nucleofection did not induce non-specific effects on gene expression. Nucleofection was performed twice with a 24 hours interval. 48 h after the second nucleofection, the *NRAS* and *PTPN11* mRNA expression was analyzed by Q-PCR (*GUSB* was employed as the reference gene). Cell proliferation was analyzed 0, 2 ,4 and 6 days after two repetitive transfections. Transfection efficiency was determined by flow cytometry using the BLOCK IT Fluorescent Oligo (Invitrogen Life Technologies, Paisley, UK).

#### Section 3. Cell proliferation assay

Cell proliferation was analyzed using the CellTiter 96 Aqueous One Solution Cell Proliferation Assay (Promega, Madison, W). This is a colorimetric method for determining the number of viable cells in proliferation. For the assay, 100 µL of nucleofected cells were plated in 96 wells plates 0, 2, 4 and 6 days after the last nucleofection. Plates with suspension cells were centrifuged at 800 g for 10 minutes and medium was removed. Then, cells were incubated with 100 µL/well of medium and 20 µL/well of CellTiter 96 Aqueous One Solution reagent. The plates were incubated for 1-4 hours, depending on the cell line at 37 ºC in a humidified, 5 % CO2 atmosphere. The absorbance was recorded at 490 nm using 96-well plate readers until absorbance of control cells without treatment was around 0.8. The background absorbance was measured in wells with only cell line medium and solution reagent. First, the average of the absorbance from the control wells was subtracted from all other absorbance values. Data were calculated as the percentage of total absorbance of siRNA transfected cells/absorbance of control cells.

#### Section 4. Quantitative-PCR (Q-PCR)

The expression of *NRAS* and *PTPN11* was analyzed by Q-PCR in HL-60, HEL, MV4-11 and OCI-AML3 AML cell lines. First, total mRNA was extracted with Trizol© Reagent 5791 (Life Technologies, Carlsbad, CA, USA) following the manufacturer instructions. RNA concentration was quantified using NanoDrop Specthophotometer (NanoDrop Technologies, USA). cDNA was synthesized from 1 µg of total RNA using the PrimeScript RT reagent kit (Perfect Real Time) (Cat No RR037A, TaKaRa) following the manufacturer's instructions. The quality of cDNA was checked by a multiplex PCR that amplifies *PBGD*, *ABL*, *BCR* and *β2-MG* genes. Q-PCR was performed in a QuantStudio 5 Real-Time PCR System (Applied Biosystems), using 20 ng of cDNA in 2 µL, 1 µL of each primer at 5µM (NRAS F: 5'-CGCACTGACAATCCAGCTAA-3'; NRAS R: 5'-CCAACAAACAGGTTTCACCA-3'; PTPN11 F: 5'-CGGAGCCTGAGCAAGGAG-3'; PTPN11 R: 5'-CTGCCTCCACACCAGTGATA-3'; GUSB F:5' gaaaatatgtggttggagagctcatt-3'; GUSB R: 5'-ccgagtgaagatccccttttta-3'), 6 µL of SYBR Green PCR Master Mix 2X (Cat No 4334973, Applied Biosystems) in 12 µL reaction volume. The following program conditions were applied for Q-RT-PCR running: 50 ºC for 2 min, 95 ºC for 60 s following by 45 cycles at 95 °C for 15 s and 60 °C for 60 s; melting program, one cycle at 95 °C for 15 s, 40 °C for 60 s and 95 °C for 15 s. The relative expression of each gene was quantified by the Log $2^{(-\Delta\Delta Ct)}$ method using the gene *GUSB* as an endogenous control.

#### Section 5. Statistical pipeline

A statistical pipeline (**Figure S1**) has been developed to solve the problem of identifying subrogate mutation biomarkers of gene essentiality (RNAi target genes).

Data of RNAi libraries (more than 17,000 knocked-down genes in 412 cancer cell lines) of the project Achilles [1] were integrated with their corresponding mutational profiles (mutations in ~1600 genes; Figure 1A) obtained from the Cancer Cell Line Encyclopedia (CCLE)[2] and Shao et al. [3]. We filter out the mutations that meet any of the following criteria: (i) common polymorphisms, (ii) allelic fraction < 10%, (iii) putative neutral variants (missenses present in less than 2 warm-blooded vertebrates), or (iv) located outside of the coding sequence. We used the DEMETER score [4,5] as a measure of gene essentiality of the RNAi libraries of the project Achilles [1]. DEMETER quantizes the competitive proliferation of the cell lines controlling the effect of off-target hybridizations of siRNAs by solving a complex optimization problem. The more negative the DEMETER score is, the more essential the gene is for a cell line. We imputed missing elements of DEMETER using the nearest neighbor averaging algorithm [6]. In addition, we collected gene expression patterns from RNA-seq data [7] to confirm that essential genes are expressed when they are essential.

Based on DEMETER data, we first identified genes that were essential for a selected tumor subtype. Essential genes were required to meet several criteria: i) they must be essential for at least 20% samples of the selected cancer subtype, ii) they must be specific to the cancer type under study, i.e. they must be non-essential for other cancer types and iii) they must be expressed before RNAi experiment (>1TPM at least in 75% samples). These filters reduce the number of hypotheses in the statistical analysis.

We developed a statistical algorithm to identify genes whose essentiality is highly associated with the mutational status of other genes. Dealing with this statistical issue implies solving a large multiple hypotheses problem (more than one million hypotheses). In similar scenarios, traditional corrections -such as Benjamini-Hochberg (BH), Bonferroni or Holm- showed very few or no gene-biomarker pairs for a given FDR [8]. In order to overcome this problem, we developed a covariate-based statistical approach -similar to the Independent Hypothesis Weighting procedure [8].

Statistical model: Let $e$ denote the number of RNAi target genes and $n$ denote the number of screened samples. Let **D** be an $e \times n$ matrix of essentiality whose entries $d_{ij}$ represent the

DEMETER score for the RNAi target *i* in sample *j*. Let **m** be a $m \times n$ dichotomized matrix whose entry $m_{ij}$ denotes whether sample *j* is mutant or not according the previous criteria:

$$m_{ij} = \begin{cases} 1, & if mutant(MUT) \\ 0, & if\ wild - type\ \ (\mathrm{WT}) \end{cases}, \quad (1)$$

Let **s** be a subset of n' cell lines that yields an essentiality vector$\boldsymbol{d_s} = \left(d_{e_{s_1}}, \ldots, d_{e_{s_{n'}}}\right)$ for the e[th] RNAi target. Let $\boldsymbol{m_s} = \left(m_{s_1} \ldots, m_{s_n}\right)$ be the expression vector of a putative gene biomarker. The null hypotheses are defined as:

$$H_0^g: E(\boldsymbol{d_s}|\boldsymbol{m_s} \in MUT) = E(\boldsymbol{d_s}|\boldsymbol{m_s} \in WT) \quad (2)$$

This null hypothesis is therefore: "the expected essentiality of a gene knock-down is identical in mutant and wild-type cell lines". To test this hypothesis, we used a moderated t-test implemented in *limma*[9]. We applied this test for each RNAi target and all the mutations to get the corresponding p-values. Dealing with these p-values implies correcting for multiple hypotheses.

In order to face these challenges, we followed a methodology similar to the IHW (Independent Hypothesis Weighting) procedure [8], which increases the power of a test by grouping the results using covariates. We show in the main manuscript that the number of positives returned by IHW is larger for all the cancer datasets and therefore, this method outperforms the standard FDR estimation both in sensitivity and specificity (as shown in the lemma of the following section in the supplementary material).

In our case, we divided the p-values corresponding to all the tests into 2n groups, where n is the number of knock-down genes.

For each of these groups, we computed the local false discovery rate (local fdr) [10]. The local fdr estimates, for each test, the probability of the null hypothesis to be true, conditioned on the observed p-values. The formula of the local fdr is the following:

$$P(H_0|z) = localfdr(z) = \frac{\pi_0 f_0(z)}{f(z)}, \quad (3)$$

where *z* are the observed p-values, $\pi_0$ is the proportion of true null hypotheses –estimated from the data-, $f_0(z)$ the empirical null distribution –usually a uniform (0,1) distribution for well-designed tests- and $f(z)$ the mixture of the densities of the null and alternative hypothesis, also estimated from the data.

As stated in [47], "the advantage of the local fdr is its specificity: it provides a measure of belief in gene i's 'significance' that depends on its p-value, not on its inclusion in a larger set of possible values" as it occurs, for example with q-values or the standard False Discovery Rate. The local fdr and $\pi_0$ were estimated using the Bioconductor's R Package *qvalue* [11].

For a selected cohort of cells, the algorithm outputs a ranking of significant gene pairs (GPs) that consist of a couple of genes in which the first one is essential depending on the mutational status of the other. For the final ranking, we selected those GPs that showed a p-value < 0.05 and local FDR ≤ 0.6, |ΔDEMETER| > 2. Additionally, we interrogated which of these pairs had direct relationships (co-expressed, annotated in the same pathway database or contained in a common experiment) in the STRING database [12] to ensure there is an established biological relationship between the essential gene and the subrogate biomarker. This biological double-check is not necessary and can be omitted when the researcher looks for novel relationships.

#### Section 6. A larger number of positives outperforms specificity and sensitivity

**Lemma:** *Let us consider two methods that correct multiple hypothesis test, and let us consider that both methods provide a different number of positives for the same FDR. Then, the method that provides a larger number of positives has more statistical power. It is also more specific and sensitive.*

The power or sensitivity of a statistical test is the probability that the test correctly rejects the null hypothesis $H_0$ when the alternative hypothesis $H_1$ is true. Its value is *TP/(TP+FN)*.

Let's consider that the estimation of the FDR is performed by two tests A and B and both tests have the same False Discovery Rate (20% for example). The FDR will be

$$FDR = \frac{FP_A}{TP_A + FP_A} = 1 - \frac{TP_A}{TP_A + FP_A} = \frac{FP_B}{TP_B + FP_B} = 1 - \frac{TP_B}{TP_B + FP_B} (1a)$$

The power of each test will be

$$PW_A = 1 - \beta_A = \frac{TP_A}{TP_A + FN_A} (1b)$$

$$PW_B = 1 - \beta_B = \frac{TP_B}{TP_B + FN_B} (1c)$$

Since both tests are performed on the same dataset, the number of true null hypothesis $H_0$ (FP + TN) and true alternative hypothesis $H_1$ (TP+FN) will be identical, i.e.,

$$FP_A + TN_A = FP_B + TN_B (1d)$$

$$TP_A + FN_A = TP_B + FN_B (1e)$$

Notice that the denominators of the expression of the power (eq (1b) and (1c)) are identical according to (1e).

The total number of positives returned by each test is $TP_A + FP_A$ and $TP_B + FP_B$. Let's assume that method A, returns more positives than method B, i.e.

$$TP_A + FP_A > TP_B + FP_B (2)$$

Using eq. (1a), and (2)

$$TP_A = (1 - FDR)(TP_A + FP_A)(3a)$$

And,

$$TP_B = (1 - FDR)(TP_B + FP_B)(3b)$$

Since (2), the righthand member of equation (3a) is larger than the righthand member of equation (3b) and therefore,

$$TP_A > TP_B (4)$$

As a result,

$$PW_A > PW_B \blacksquare$$

Corolary I. Since $PW_A = 1 - \beta_A$ the type II error using A is smaller than using B.

$$\beta_A < \beta_B$$

Corolary II. The type I error is

$$\alpha_A = \frac{FP_A}{FP_A + TN_A}$$

And the sensitivity is:

$$1 - \alpha_A = \frac{TN_A}{FP_A + TN_A}$$

By (1e) and (4), it is straightforward to conclude that

$$\alpha_A < \alpha_B$$

Therefore, the method that provides a larger number of positives outperforms the other both in terms of specificity and sensitivity (or type I and type II errors).

#### Section 7. Comparison between CRIPSR and RNAi experiments

Some of the cell-lines in CCLE has been interrogated using both RNAi and *CRISPR-Cas9* experiments. This overlap opens the possibility to investigate if the results from both experiments are coherent.

We compared the results of CRIPSR (using CERES to deconvolve the signal) and RNAi (using DEMETER and DEMETER2) to deconvolve the signals and get the final scores.
There were 140 common cell-lines and 14189 common genes with the corresponding annotated mutations. Each of the scores (CRISPR, DEMETER2 and DEMETER) were stored as matrices (genes in rows and cell-lines in columns).

The first analysis performed was to compute the correlation of the scores for each cell-line vs the others, i.e. compute the column-wise correlation matrix. This matrix is expected to have a mean value well above zero since many essential genes are likely to be essential for a large number of cell-lines and many non-essential genes are expected to be non-essential for other cell-lines. This is the case for CRISPR (average correlation around 0.8), DEMETER2 (average correlation around 0.7) but not for DEMETER (average correlation around 0) (see Figure S27).

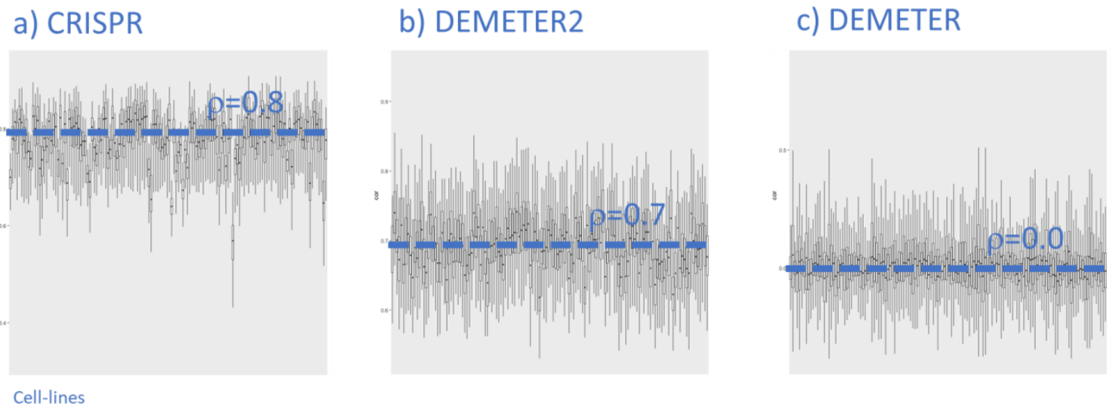

*Figure S27. Comparison of the correlation of essentiality scores across cell-lines. Each boxplot represents the correlation between a particular cell-line and all the others (excluding the cell-line itself). The y-axis are different in the three panels. A large correlation is expected in this case, since most essential genes are expected to be essential for a large number of cell-lines and most non-essential genes are expected to be non-essential for a number of cell-lines.*

We also computed the correlation of the scores for each gene and compute the density function of these correlations (in this case, we computed the row-wise correlation matrix comparing each combination of two of the three methods). DEMETER and DEMETER2 have a very strong correlation (average value around 0.9). CRISPR and DEMETER or DEMETER2 have a weaker correlation (ro = 0.03). Despite this low value, this correlation is strongly significant (one-sample t-test p-value < 2.2 e-16) (Figure S28).

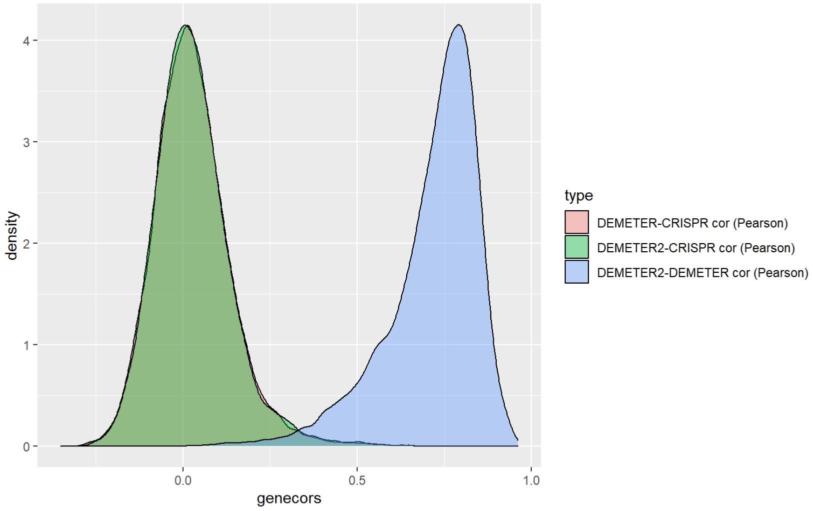

*Figure S28. Density function of the correlation between the scores across genes.*

A possible interpretation of these results is the following. DEMETER and DEMETER2 provide very similar scores (as shown by the strong correlation across the genes). The main difference between them is the different offset of these values across the genes.. This offset in DEMETER2 seems to improve the results since the correlation of the scores across the cell-lines is well above zero as shown in Figure S27.

The (extremely) weak albeit significant correlation between CRISPR and DEMETER (or DEMETER2) drove us to study more in depth which is the relationship between both scores.

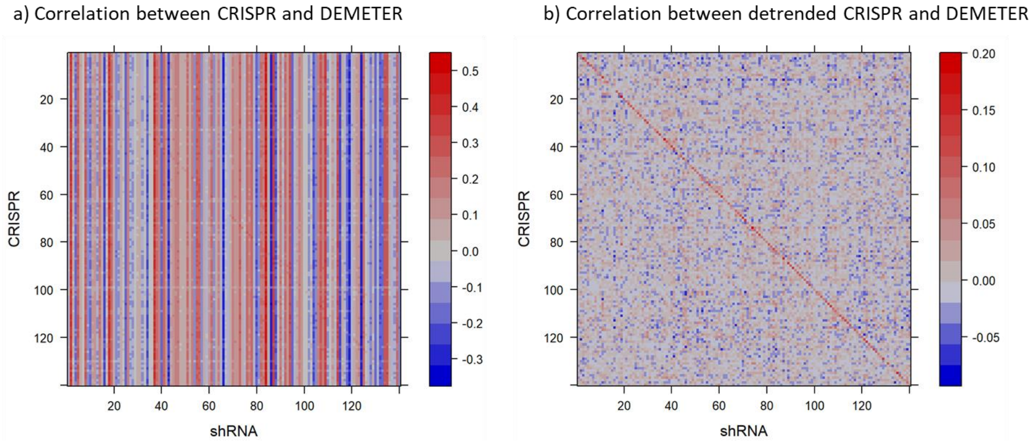

*Figure S29. Correlation between CRISPR and DEMETER. Panel a): Original correlation. Panel b): Correlation after removing the first principal component of the correlation.*

Panel "a" in Figure S29 shows the correlation between CRISPR and DEMETER across the cell-lines. This correlation shows distinctive vertical bars: if the scores of a cell-line have a strong correlation with another cell-line, then they are also highly correlated with all the others. This fact made us think that the offset for each gene and for each cell-line was the main confounding factor. We removed this offset with the aid of the singular value decomposition. After removing the first component, panel "b" of Figure S29 shows that the correlation matrix of the expression of the cell-lines is closer to the identity matrix (a relatively large positive value in the diagonal, and values close to zero in the off-diagonal entries). As a result, CRISPR and RNAi experiments are coherent, but a special attention must be taken to the offset, i.e. it is better to base any analysis on incremental values instead of absolute values as we are doing in the main manuscript.

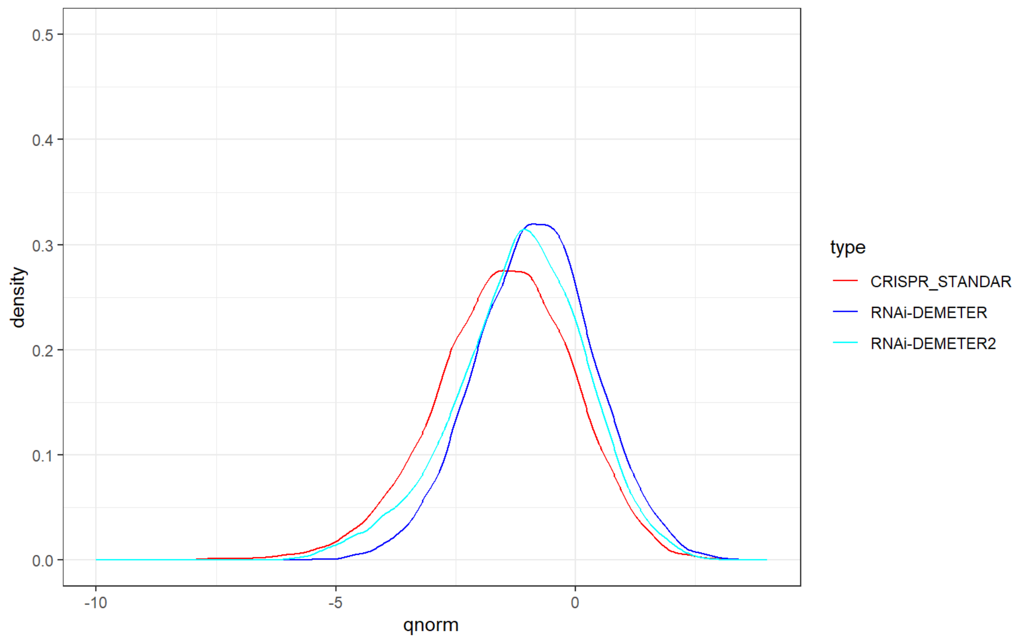

*Figure S30. Comparison between CRIPSR, DEMETER and DEMETER2 based on how they reflect the tissue of origin.*

Another question that arises is "which one -RNAi or CRISPR- better reflects the tissue of origin?". The overall idea is that if the scores for RNAi or CRISPR were dominated by noise they would not show any relationship with the tissue of origin. If either of the techniques is coherent with the tissue of origin, the signal to noise ratio of the technique is likely to be better than if no relationship appears.

We build a linear model in which the design matrix reflects the tissue of origin. Using limma we solved for each gene and compute the F-test of an ANOVA type I comparing the tissue of origin. The p-values of these tests were significant in most of the cases.

Figure S30 shows the density function of the z-scores that corresponds to each of these p-values. The more negative the z-score is, the more related are the scores with the tissue of origin (negative values are better). DEMETER and DEMETER2 are very close to each other, reinforcing the fact that the main difference between them is a more appropriate selection of the offset in DEMETER2. Nevertheless, since the density functions are not identical, there also other differences in the results that -at least in this comparison- make DEMETER2 a better method than DEMETER. CRISPR outperform both of them but not by a large margin.
